# An oral protein cocktail therapeutic for *C. difficile* infection

**DOI:** 10.1101/2021.12.21.473715

**Authors:** Hui Zhao, Michael Dodds, Alex Pollock, Mesfin Gewe, Anissa Martinez, Kristie Keeney, Benjamin W. Jester, Melanie Hutton, Hannah Tabakh, Mia Zhang, Mark Heinnickel, Troy Paddock, Stacey Ertel, Kanchan Aggarwal, Caroline Amendola, Michael Tasch, Colin Brady, Nathaniel R. Sanjaya, Madison Kahwaty, Kendra Cruickshank, Sarah Struyvenberg, Jason Dang, Tracy Saveria, Chelsea Shanitta, Yena Park, David Doughty, Jackie Poirot, Caitlin Pensak, Thomas Adame, Jeremy Ferrara, Aaron Read, Dalton Heil, Spencer Shibuya, James “Chip” Henderson, David Fletcher, Taylor Charron, Kristjan Sigmar, Eva Aw, René Ruiz, Karen Yoshino, James Lo, Dorothy Strong, Lauren Goetsch, Caitlin Gamble, Sarah Shephard, Yasi Zhong, Ryan Antolock, Cosette Loper, Gabby Guzman, Sophie Gist, Asa Davis, Sanika Khare, Ashley Zambrano, Steven J. Mileto, Ryan Heselpoth, David Saunders, George B. McDonald, Dena Lyras, Vincent Fischetti, Summer Radler, Carl J. Mason, Craig A. Behnke, Nhi Khuong, Brian Finrow, James M. Roberts

**Affiliations:** Lumen Bioscience, Seattle, WA; Biomedicine Discovery Institute and Department of Microbiology, Monash University, Victoria, Australia; Laboratory of Bacterial Pathogenesis and Immunity, Rockefeller University, New York, NY

## Abstract

Here we show that transgenic spirulina may solve key challenges of developing effective monoclonal protein therapeutics for diseases afflicting gastrointestinal (GI) tissues. We describe, as a paradigm for this novel approach, LMN-201: an investigational four-component treatment for Clostridioides difficile (C. difficile) infection comprising four monoclonal protein therapeutics produced and delivered within spirulina biomass. Three antibody-like moieties within LMN-201 were rationally designed using a multiplicative potency framework, termed synthetic avidity, that increased in vitro toxin neutralization potency by >200-fold relative to the individual components. The fourth moiety, a lysin enzyme protein, adds direct, toxinotype-agnostic antibacterial activity. In rodent challenge models, species-relevant LMN-201-like cocktails reduced disease burden and mortality. In a human GI pharmacokinetic study, LMN-201 was present at concentrations 100- to 1,000-fold above the estimated minimally effective therapeutic threshold at the distal intestine. In Part A of a Phase 2 study, participants with C. difficile infection (CDI) were treated for seven days with LMN-201 plus standard-of-care antibiotics. The primary endpoints of safety and tolerability were assessed favorably. An interim analysis of efficacy in pre-planned coordination with the data and safety monitoring board showed an initial clinical cure rate of 21/21 (100%) and sustained clinical cure rate through four weeks of 19/21 (90.5%). These findings support LMN-201 with standard-of-care antibiotics as a viable treatment option for CDI, validate spirulina as a scalable platform for oral biologics, and establish a generalizable strategy for engineering high-potency combination protein therapeutics for diseases with a GI tissue nexus.

## Introduction

The most potent biologic drugs—especially protein-based therapies—have highly specific cellular or molecular targets yet are typically delivered throughout the body by infusion into the bloodstream. This reduces efficacy in many cases due to limited partitioning into the intended tissue compartment and increases the risk of off-target toxicities. Nevertheless, for drugs directed at gastrointestinal (GI) targets the default often continues to be systemic delivery. The GI tract rapidly degrades dietary protein and the same would be expected of ingested protein drugs, leading to a substantial reduction of therapeutic effect. Indeed, with few exceptions (e.g., enzyme supplements), there are no U.S. Food and Drug Administration (FDA)-approved, orally delivered protein biologics intentionally designed to achieve therapeutic activity in the GI lumen. However, in healthy individuals the concentration of serum proteins like immunoglobulin G (IgG) is 1,000- to 10,000-fold greater in the blood than in whole GI lavage fluid^1^, consistent with minimal baseline partitioning into the GI lumen. Even in individuals with epithelial barrier disruption, such as those with severe inflammatory bowel disease, GI luminal concentrations of systemically delivered antibodies are modest and variable^2,3^. It is therefore especially challenging to achieve therapeutic GI luminal drug concentrations by injection.

Clostridioides difficile (C. difficile) is an opportunistic GI pathogen that causes severe, recurrent diarrhea and colitis and is associated with significant morbidity and mortality worldwide^4,5^. It is the most common cause of healthcare-associated infections in U.S. hospitals, where nearly half a million cases occur each year^5^. C. difficile virulence is mediated by two exotoxins, TcdA and TcdB^6–9^. The toxins initially engage cell surface receptors on intestinal epithelial cells then translocate to the cytoplasm where they disaggregate the actin cytoskeleton resulting in disruption to the colonic epithelium^10–12^. Current evidence indicates that TcdB is the principal driver of human disease^8,13–18^.

Antibiotic therapy initially cures approximately 80% of C. difficile infections (CDIs). However, 20–40% of patients suffer recurrence within weeks of initial cure^19^ and the chance of additional CDI recurrence in these patients exceeds 40%^20,21^. The principal cause of CDI is thought to be microbial dysbiosis caused by antibiotics, which creates a niche favorable to C. difficile proliferation^22^.

A major medical need could be addressed with an antimicrobial therapy that is more effective in achieving initial cure and more specific to the C. difficile pathogen, thereby decreasing disruption of the normal microbiome and reducing the likelihood of recurrence. Bezlotoxumab, a monoclonal antibody against TcdB, failed to improve CDI primary cure rates or mortality, but was approved by the FDA for preventing CDI recurrence^18,23^. However, like other systemically delivered biologics used to treat GI diseases, it suffered from poor efficacy and significant off-target toxicities, including a risk of heart failure noted on the label^18,24^, and it was discontinued by the manufacturer so is unavailable in the U.S. market. Fecal microbiota transplant products—which cannot be administered in combination with standard-of-care (SoC) antibiotics and which do not directly target the pathogen—are now the only approved available therapeutic option for reducing CDI recurrence^25,26^.

The recent discovery of methods for genetically engineering spirulina—an edible photosynthetic alga that has historically resisted genetic manipulation—allowed its rapid development as a new platform for the manufacture and oral delivery of protein biologic drugs^27^. Spirulina expresses exogenous protein drugs intracellularly in amounts that can exceed 10 percent of biomass, and high production yields can be achieved with indoor cultivation in photobioreactors equipped with LED lights as the sole energy source and carbon dioxide as the sole carbon source. Processing the harvested spirulina biomass in a conventional spray dryer is the only downstream step needed for preparation of the final drug product: the protein drug bioencapsulated within whole desiccated, inviable spirulina biomass.

The key advance reported here is how the spirulina biomanufacturing and drug delivery platform addresses the challenges of orally delivering protein drugs to targets within GI tissues. We highlight manufacturability: the high expressivity of exogenous proteins combined with minimal downstream processing required for oral delivery makes it practical to administer clinically relevant doses of the drug product, if necessary, multiple times per day. We also highlight how spirulina biomanufacturing facilitates assembly of biologic drug cocktails, which can have potencies that exceed those possible with monotherapies. Higher potency increases the potential for therapeutic efficacy at lower intraluminal concentrations, which may be critical given the challenges of oral delivery.

Following preclinical studies using rodent models of CDI, we evaluated the effectiveness of this approach in humans by first assessing the intraluminal pharmacokinetics of the investigational spirulina-assembled oral drug cocktail in a Phase 1 clinical trial. We found that the combination of high expressivity and high potency achieved concentrations of the biologic drug cocktail in the distal small intestine that met or exceeded estimated thresholds for therapeutic effectiveness. A subsequent Phase 2 trial showed that twice-daily administration of LMN-201 for seven days alongside SoC antibiotics both increased the CDI initial cure rate and reduced four-week recurrence relative to historical data for both antibiotics alone and antibiotics in combination with bezlotoxumab.

In sum, this approach offers a generalizable strategy for luminal delivery of monoclonal protein therapeutics, potentially addressing unmet needs across a wide range of diseases and disorders linked to GI tissues or physically present in the GI lumen itself.

## Results

### Development and characterization of the spirulina-engineered anti-C. difficile therapeutic cocktail, LMN-201

LMN-201 was designed as a four-component therapeutic cocktail whose active moieties are four monoclonal proteins: a catalytic enzyme derived from PlyCD, a C. difficile-specific endolysin^28^, and three TcdB-binding proteins derived from camelid single-domain antibody fragments (VHHs) previously shown to inactivate TcdB^9,29–31^. The catalytic enzyme derived from PlyCD, PlyCD (protein product: pp1092), was encoded with a C-terminal hexahistidine (6×His) tag. The three VHH-containing proteins, pp1005, pp1006, and pp1007, are self-assembling homodimers derived from the VHHs 5D, E3, and 7F, respectively, linked with a 5HVZ dimerization domain, a maltose-binding protein chaperone, and a C-terminal 6×His tag (Figure 1A).

**Figure 1:**
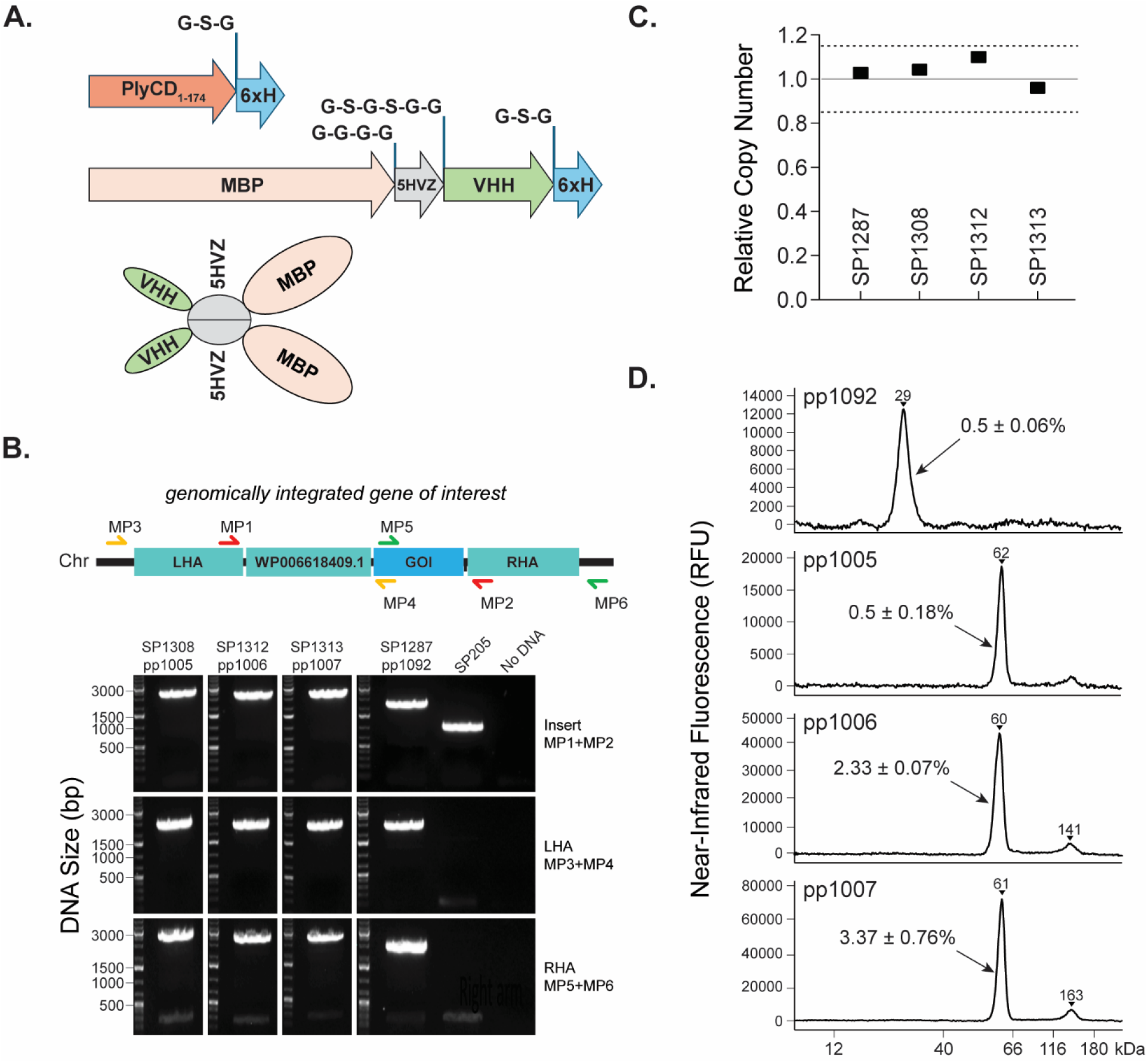
Spirulina Strain Engineering and Characterization. **(A)** Schematic of modified PlyCD_1–174_ (pp1092) and TcdB-binding proteins (pp1005, pp1006, pp1007). **(B)** Schematic of the genomic integration locus and PCR-based analysis of integration specificity and segregation. PCR using primers MP1/MP2 produced bands of the expected sizes in all engineered strains, with no detectable parental alleles, indicating complete segregation to homozygosity. Primer pairs used to amplify the left homology arm **(LHA)** and right homology arm **(RHA)** each included one primer (MP3 and MP6, respectively) that anneals uniquely to the spirulina genome at the target locus and is absent from the donor plasmid. PCR amplification of the LHA and RHA using MP3/MP4 and MP5/MP6, respectively, confirmed site-specific genomic integration. All PCR products were sequence-verified and matched the expected designs. **(C)** Relative copy numbers of genes of interest (GOIs) determined by next-generation sequencing, calculated by comparing read counts for each gene of interest with those of ten randomly selected native spirulina genes. The dotted line indicates the range within which relative copy numbers were considered equivalent to one. **(D)** Capillary electrophoresis–based immunoassay chromatograms of pp1092, pp1005, pp1006, and pp1007 extracted from spirulina biomass and analyzed under reducing conditions. Soluble protein abundance, expressed as a percentage of dry biomass, is indicated in each panel.

The gene encoding each protein was separately integrated into the genomes of spirulina strain SP205 by homologous recombination, and a single clonal filament was isolated from each transformation (Figure 1B). The final spirulina strains (SP1287, SP1308, SP1312, and SP1313) were confirmed homozygous with respect to the relevant transgenes (Figure 1B, 1C) and were then used to establish a frozen strain bank. After spray drying, pp1092, pp1005, pp1006, and pp1007 accounted for 0.5%, 0.5%, 2.3%, and 3.4% of the total dried biomass of their respective spirulina strains (Figure 1D).

Expressed and purified proteins were confirmed to exist in their designed monomeric (pp1092) or dimeric (pp1005, pp1006, pp1007) states using size-exclusion chromatography and electrophoresis in reducing and non-reducing conditions (Figures 2A, 2B). Intra-scaffold disulfide bonds were present in the extracted proteins (Figure 2B) but were dispensable for stable assembly of the 5HVZ-VHH dimers (Figure 2C). Protein pp1092 isolated from SP1287 was confirmed to have lytic activity against C. difficile strain ATCC 43255; the potency was not substantially different from the same protein expressed in E. coli (Supplemental Figure 4). An analysis of binding kinetics using biolayer interferometry indicated apparent equilibrium dissociation constants (k_D_) in the mid- to low-picomolar range for dimeric versions of all three VHHs, reflecting substantially greater apparent avidity than that observed for monomeric versions (Figure 2D).

**Figure 2:**
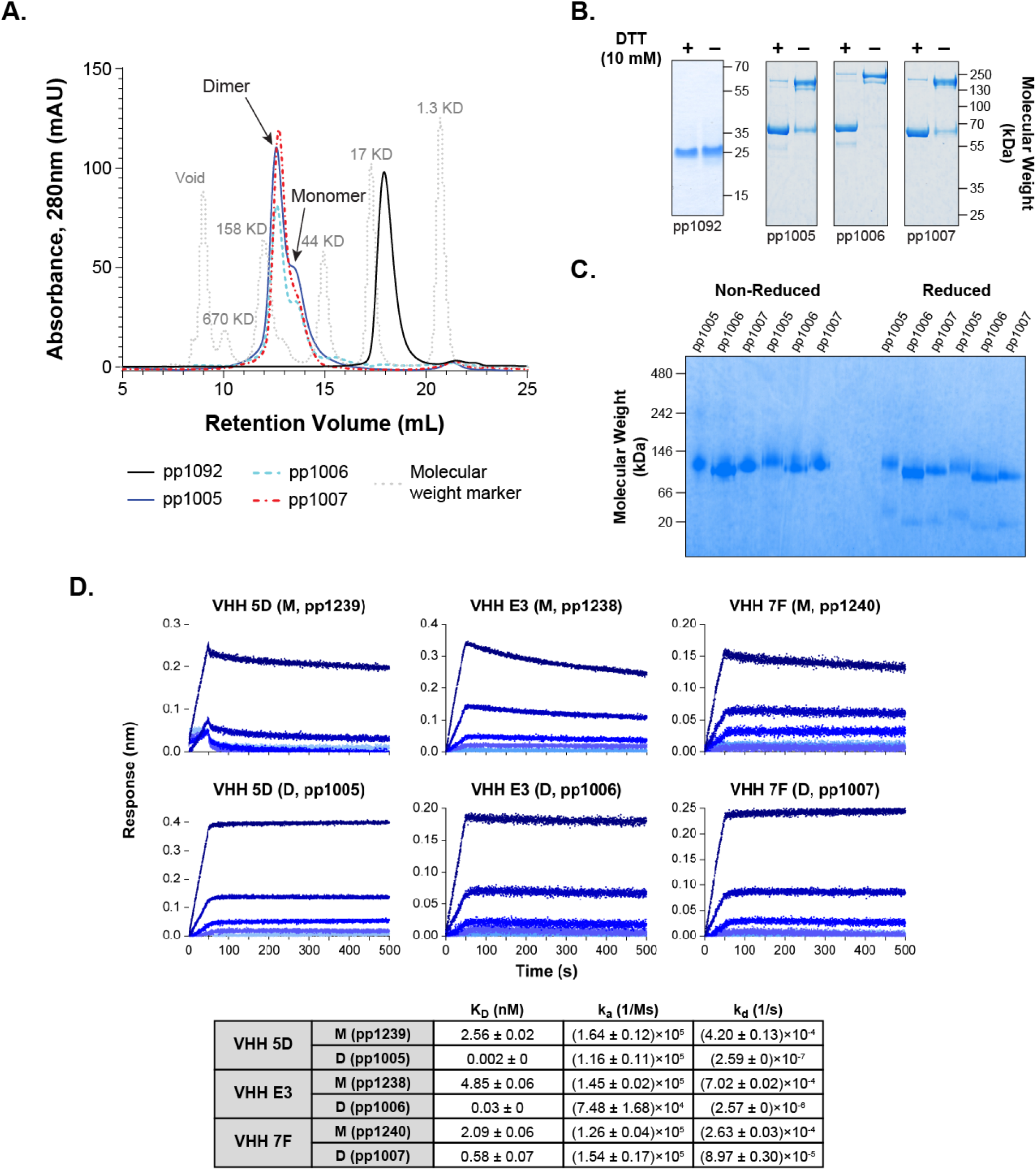
Characterization of Spirulina-expressed TcdB-binding Proteins and Lysin. **(A)** Size-exclusion chromatography profiles of spirulina-expressed and purified pp1092, pp1005, pp1006 and pp1007 in aqueous solution (PBS, pH 7.4), analyzed on a Superdex 200 10/300 GL FPLC column. Grey traces indicate molecular mass standards. Elution peaks corresponding to monomeric and dimeric species are indicated. **(B)** Assessment of inter-monomer disulfide bond formation by denaturing gel electrophoresis under non-reducing and reducing (10 mM DTT) conditions for pp1092, pp1005, pp1006, and pp1007. (C) Native gel electrophoresis of pp1005, pp1006, and pp1007 under non-reducing and reducing (50 mM DTT) conditions. The dimeric state of the TcdB-binding proteins was largely preserved in the presence of reducing agent. (D) Biolayer interferometry sensorgrams showing binding kinetics of anti-TcdB VHHs 5D, E3 and 7F expressed as monomers (pp1239, pp1238 and pp1240) or dimers (pp1005, pp1006 and pp1007). Association (**k⍰**) and dissociation (**k_d_**) rate constants were derived from concentration-dependent binding responses, and apparent equilibrium dissociation constants (K_D_) were calculated from kU and k_d_. Kinetic parameters are summarized in the table (**M**, monomer; **D**, dimer).

### Multiplicative potency for in vitro toxin neutralization

We used a cell-based toxin-neutralization assay to quantify potency of spirulina-expressed TcdB-binding proteins in different concentrations and combinations against four clinically important TcdB types (1.1, 2.1, 3.1, and 5.1) across a wide range of clinically relevant concentrations^32^. TcdB1 toxicity data were collected and fit to a four-parameter sigmoid using nonlinear regression to minimize the sums of residuals squared (see Supplemental Materials; Supplemental Table 1). The concentration of half-maximal inhibition (IC_50_) of cell adhesion by TcdB1 (the most prevalent variant identified in infected patients) was 1.268 fM, with a Hill slope coefficient of 1.495 (Supplemental Table 2; Figure 3A).

**Figure 3:**
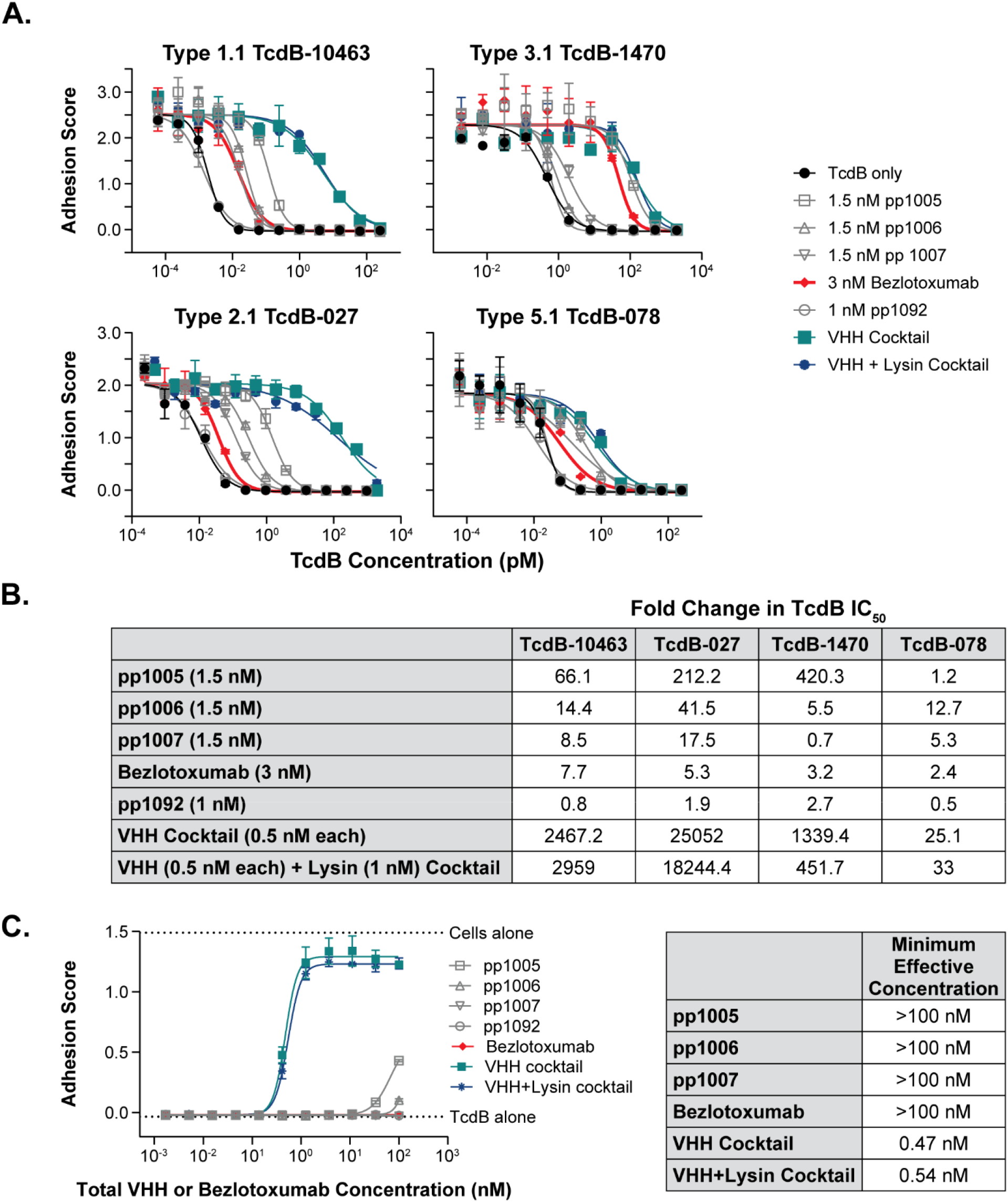
Comparative Neutralization of the Four Clinically Important TcdB Types by LMN-201 and Bezlotoxumab. **(A)** Neutralizing activity of pp1005, pp1006 and pp1007, individually or as a triple cocktail, against TcdB toxin types 1.1 (TcdB-10463), 2.1 (TcdB-027), 3.1 (TcdB-1470) and 5.1 (TcdB-078). 1.5 nM of each TcdB-binding protein or the cocktail of all three TcdB-binding proteins (0.5 nM of each) was tested against serially diluted TcdB with Vero cells on the xCELLigence system (see Methods). For comparison, bezlotoxumab was tested at 3 nM under identical conditions. To test for interference from the lysin component, the effect of 1 nM pp1092 on cocktail activity was also assessed. **(B)** Fold change in TcdB IC₅₀ values in the presence of each treatment, calculated from the data shown in A. **(C)** Neutralization of TcdB-027 at a high, clinically relevant toxin concentration. pp1005, pp1006 and pp1007 were tested individually or as a cocktail, with or without lysin. The cocktail contained each TcdB-binding protein and pp1092 at their therapeutic molar ratio (1:2.5:5.3:9.8). Individual TcdB-binding proteins, lysin, cocktails and bezlotoxumab were serially diluted and tested against 100 pM TcdB. The horizontal axis indicates the total TcdB-binding protein or bezlotoxumab concentration.

Data were collected to evaluate inhibition of TcdB toxicity by each TcdB-binding protein alone, and all two- and three-way combinations (Figure 3A; Supplemental Figure 1). An established C. difficile treatment (bezlotoxumab) was included for comparison. Each of the TcdB-binding proteins had neutralizing activity against most of the four clinically important TcdB toxin types, with sub-nanomolar K_i_values (the concentration that shifts TcdB toxicity by 2-fold) against TcdB1 (Supplemental Table 2).

The triple cocktail strongly neutralized all tested toxin types with greater-than-additive potency, providing robust neutralization across the relevant TcdB concentrations (Figure 3A). Specifically, 1.5 nM of any one of the three TcdB-binding proteins shifted the IC_50_ of Type 1.1 TcdB by a factor of 8.5–66.1, whereas a 1.5 nM cocktail (0.5 nM each) shifted the IC_50_ by a factor of more than 2,000 (Figure 3B). For Type 2.1 TcdB the potency of the cocktail was similarly striking. At a concentration of 1.5 nM, the individual TcdB-binding proteins shifted the IC_50_ by 17.5–212.2-fold, while a 1.5 nM cocktail (0.5 nM each) shifted the IC_50_ more than 25,000-fold. As a control for interference, it was shown that pp1092 did not affect toxin neutralization by the TcdB-binding-protein cocktail (Figure 3). Compared to the triple-VHH cocktail, bezlotoxumab was less potent at neutralizing all tested toxin types in vitro, and its neutralizing activity against Type 2.1 was notably weak, with a potency approximately 3,000–5,000-fold less than the TcdB-binding-protein cocktails.

The multiplicative activity of the TcdB-binding-protein cocktail was also illustrated in a more stringent in vitro test against Type 2.1 TcdB at a concentration of 100 pM, at or near the upper end of the clinically relevant range^33^ (Figure 3C). Because this TcdB concentration is four orders of magnitude greater than its IC_50_, neutralization of approximately 99.99% of the TcdB would be required to reduce cell killing by 50%. We designated the concentration required to cause this 50% reduction in cell killing by this high amount of TcdB the “minimum effective concentration” (MEC). The MEC of the triple cocktail (both with and without pp1092) under these conditions was approximately 0.5 nM. The MEC of the cocktail was at least 200-fold lower than the individual cocktail components and bezlotoxumab, all of which had MECs greater than 100 nM.

### In vitro protease sensitivity of LMN-201

The susceptibility of orally administered protein drugs to proteolytic degradation in the GI tract is likely to have a major impact on efficacy. pp1092 showed no detectable change in molecular size (25 kDa) following digestion with three major GI proteases: trypsin, chymotrypsin, and elastase (Figures 4A, 4B). Further, pp1092 retained >80% bioactivity in a cell-lysis assay following digestion with these proteases at estimated pre-meal concentrations (0.01 mg/mL), and >60% bioactivity following digestion at estimated post-meal concentrations (0.1 mg/mL) (Figure 4C).

**Figure 4:**
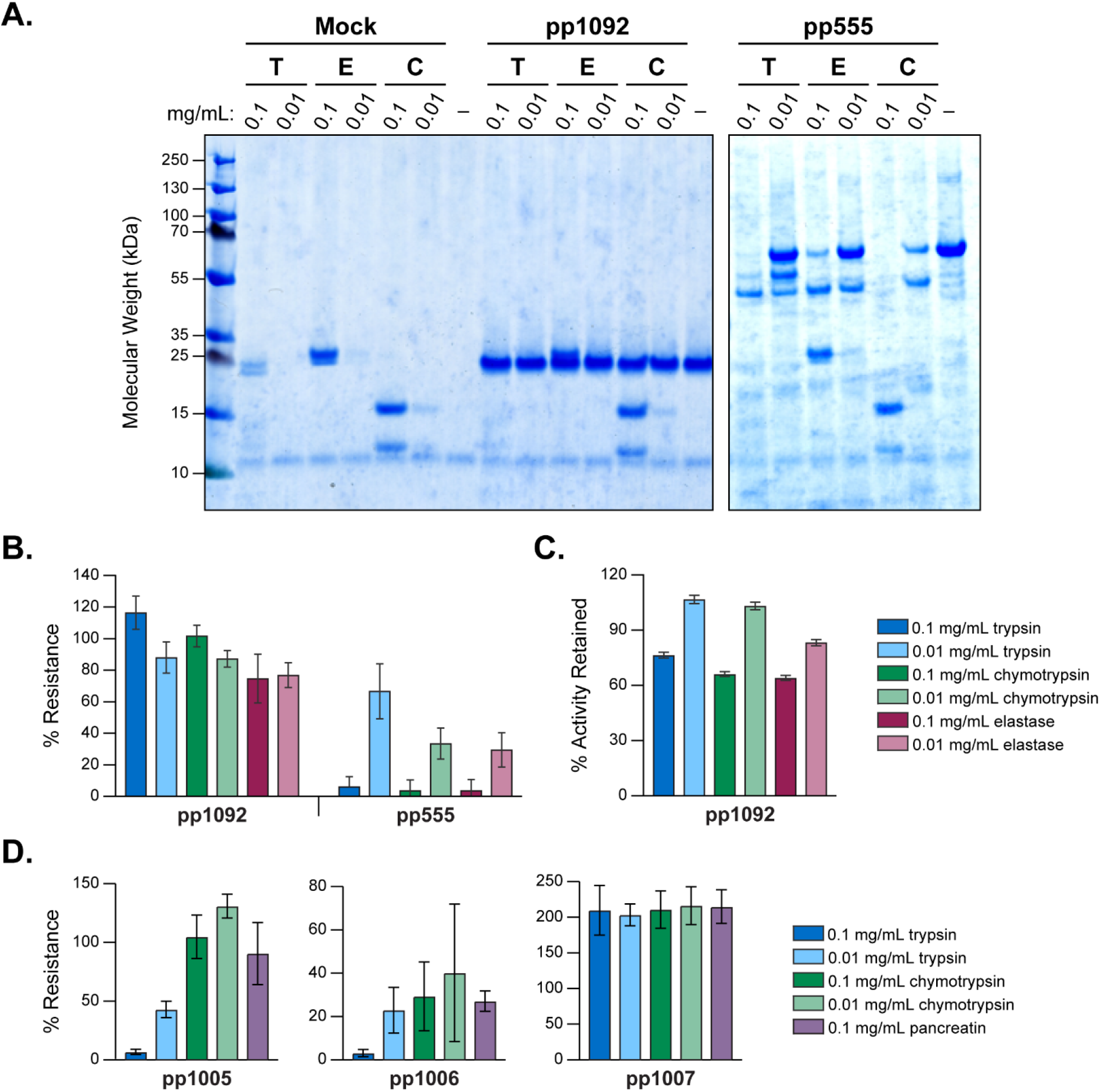
Stability of anti-C. difficile lysin and TcdB-binding proteins to gastrointestinal proteases. **(A)** Lysin PlyCD_1–174_ (pp1092, 25 kDa) and a protease-sensitive VHH (pp555, 55 kDa) were digested with trypsin (T), chymotrypsin (C), or elastase (E) at pH 6.4 for 1 hour at 37°C. Reactions were quenched with protease inhibitors, digestion products were separated by PAGE, and peptides were visualized by Coomassie staining. Mock digestions without target proteins were performed to visualize the contributions from the proteases and quenching reagents. The image shown is representative of six independent digestions. **(B)** Densitometric quantification of six independent digestions; data are presented with error expressed as SEM. **(C)** Activity of PlyCD_1–174_ following protease digestion. pp1092 was treated with the indicated concentration of protease for 1 hour at 37°C before inactivation with protease inhibitors. Residual lysin activity was measured using a lysin activity assay with Bacillus subtilis SL4 and expressed as percent activity relative to a no-protease control. Error bars represent the SEM calculated from three technical replicates. **(D)** Protease resistance profiles of the TcdB-binding proteins. Plots show TcdB-binding activity of each digested TcdB-binding protein relative to a no-protease control (% resistance). Each TcdB-binding protein was treated with the indicated concentration of protease or pancreatin for 3 hours at 37°C, followed by inactivation with a protease inhibitor. Residual binding activity was quantified by ELISA and expressed as percent resistance relative to the no-protease control. Data represent the average and standard deviation of duplicate measurements from at least three independent experiments.

At low (pre-meal) protease (trypsin, chymotrypsin, pancreatin) concentrations in simulated intestinal fluid, pp1005 and pp1006 retained 20–50% of bioactivity measured via enzyme-linked immunosorbent assay (ELISA) compared to undigested protein (Figure 4D). At higher (post-meal) concentrations, pp1005 and pp1006 exhibited significant sensitivity to trypsin, retaining <5% of their antigen-binding activities. Binding activity of pp1007 was unaffected by either protease or pancreatin. As a result, LMN-201 was administered prior to meals in clinical trials.

### In vivo preclinical efficacy of model-adapted LMN-201 analogs

To evaluate preclinical in vivo efficacy of LMN-201, we performed challenge studies in established mouse and hamster models of CDI using model-adapted cocktail formulations containing LMN-201 component proteins or closely related, species-relevant proteins.

Male C57BL/6J mice were infected with C. difficile spores and treated by oral gavage once daily with vancomycin, wild-type spirulina, Cocktail M1 (equal proportions of spirulina strains expressing pp1005, pp1006, and pp1007 and the full-length lysin PlyCD (protein product: pp1093); Supplemental Figure 4); or Cocktail M2 (same as Cocktail M1 but lacking the pp1093 lysin).

Cocktail M1 reduced fecal spore shedding by approximately two orders of magnitude compared to wild-type spirulina at 24 hours post infection (p=0.011) and at the time of euthanasia (p<0.001) (Figure 5A). Treatment with Cocktail M2 also reduced spore shedding, although the difference did not reach statistical significance at either timepoint. Maximum weight loss was significantly lower in both cocktail-treated groups compared with wild-type spirulina controls (both p<0.05; Figure 5B).

**Figure 5:**
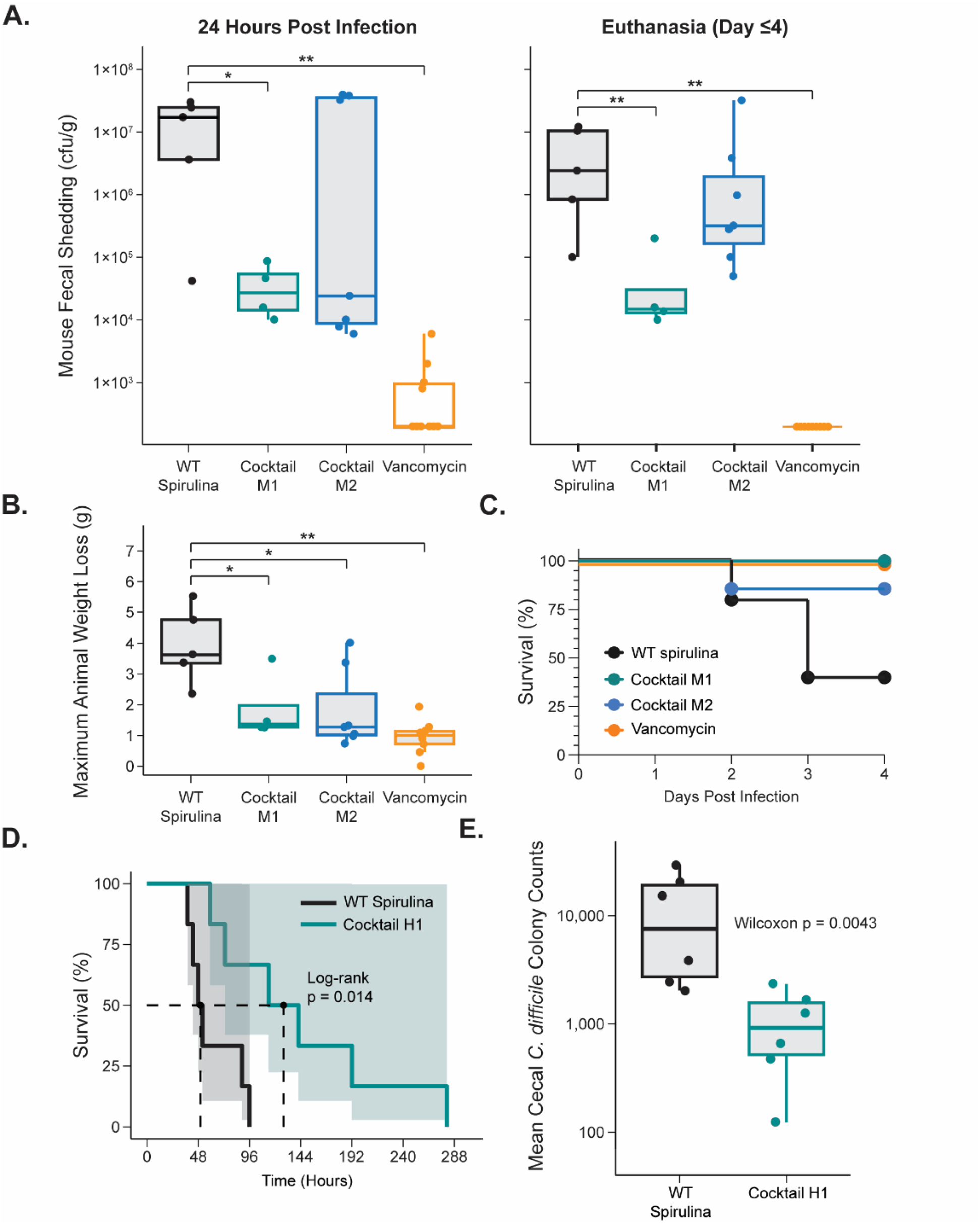
Preclinical Efficacy of LMN-201-Related Cocktails in Mouse and Hamster Challenge Models. **(A–C)** Efficacy of LMN-201 in a mouse model of Clostridioides difficile infection. **(A)** Fecal spore counts measured 24 hours post infection and at the time of euthanasia. **(B)** Maximum body-weight loss for individual animals in each treatment group. (C) Kaplan–Meier survival curves; animals were euthanized upon reaching ≥15% loss of initial body weight. In panels A and B, individual animals are shown as points, with group medians, interquartile ranges **(IQRs)**, and 1.5×IQR displayed as boxplots. **(D, E)** Efficacy of LMN-201 in a hamster challenge model. **(D)** Survival of animals treated with Cocktail H1 compared with wild-type **(WT)** spirulina controls. Median survival times were compared using the log-rank test. **(E)** Cecal C. difficile colonization in hamsters treated with Cocktail H1 compared with WT spirulina controls. *p<0.05, **p<0.001.

Survival through 4 days post infection was 100% for mice treated with vancomycin (10/10) or Cocktail M1 (4/4), 87% (6/7) for mice treated with Cocktail M2, and 40% (2/5) for mice treated with wild-type spirulina (Figure 5C). Although comparisons did not reach statistical significance, the observed trends were consistent with reductions in weight loss and bacterial shedding, supporting a potential benefit from supplementing TcdB-binding-protein cocktails with a lysin component.

In Golden Syrian hamsters, twice-daily administration of Cocktail H1 (equal proportions of spirulina strains expressing pp1005, pp1006, pp1007, pp1092 and the TcdA-binding proteins AC1 and AH3), from day –1 to day 12 significantly prolonged survival compared with wild-type spirulina controls (p=0.014; Figure 5D). Cocktail H1 also significantly reduced C. difficile colonization of the cecum relative to wild-type spirulina controls (p=0.0043; Figure 5E).

### Intraluminal pharmacokinetics of LMN-201 in participants with stable ileostomies

In a Phase 1 exploratory ileostomy study (CDI01), we measured the abundance and activities of the four therapeutic proteins in ileal fluid after oral dosing. Participants with stable ileostomies were administered LMN-201 in enteric-coated capsules. Ostomy fluid was collected hourly and evaluated for protein concentrations using quantitative bioactivity assays (see Methods). TcdB-binding proteins were detected above the minimum therapeutic concentration (MTC) for an average of 3.5–6.0 hours (Table 1). Average peak concentrations ranged from 156 to 1,787-fold above the MTC. Average total TcdB-binding protein recoveries in ileostomy outputs (relative to dosed quantities) were 1.6% for pp1005, 5% for pp1006, and 44% for pp1007.

**Table 1:**
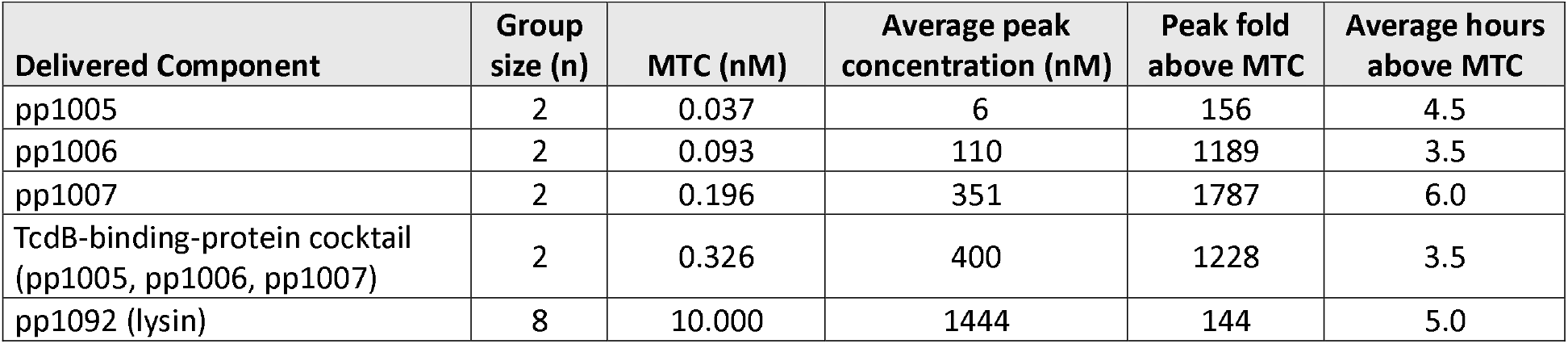
Concentrations of LMN-201 TcdB-binding Proteins and Lysin Detected in Ostomy Fluid Samples. All volunteers received pp1092 (lysin) and single (pp100#) or cocktail (pp1005, pp1006, pp1007) TcdB-binding proteins. MTC = minimum therapeutic concentration (concentration needed to reduce levels of active TcdB by 100-fold).

The MTC for pp1092 (PlyCD) was defined as the concentration of lysin required to lyse 1×10^11^ C. difficile bacteria (approximating bioburden during infection) in 1 hour^34–37^. Protein pp1092 was detected above the MTC for an average of 5.0 hours, and average peak concentrations were >100-fold higher than the MTC (Table 1). All administered treatments were safe and well tolerated; no serious or significant adverse events were reported during or after the trial.

### Manufacture of a blended cocktail for human delivery

For manufacture and clinical application, a second generation of strains was engineered to faithfully replicate the first-generation strains. To ensure consistent product potency, assays were developed for both drug substance and drug product lot release.

Lysin potency in LMN-201 was quantified using a turbidity-reduction assay (see Methods). The acceptance criterion for percent active pp1092 in spray-dried SP1287 biomass was 0.34% ± 0.08%. As expected, this potency measurement corresponded closely with the concentration determined by the capillary electrophoresis-based immunoassay, indicating that a majority of the protein was active (Figure 1D).

TcdB-binding proteins in LMN-201 were quantified using a Meso Scale Discovery (MSD) electrochemiluminescence immunoassay (see Methods). This platform enables highly sensitive, accurate, and precise multiplexed measurement of antibody-antigen interactions across a broad linear range. The acceptance threshold for percent active TcdB-binding protein in the spray-dried biomass powder of each strain was 0.4% ± 0.2% (pp1005, SP1308), 0.9% ± 0.4% (pp1006, SP1312), and 1.8% ± 0.5% (pp1007, SP1313).

If the TcdB-binding proteins’ effects on cocktail potency were linear with concentration, the K_i_ values would suggest an optimal cocktail ratio of 0.3:0.8:1 (pp1005:pp1006:pp1007). Modeling of TcdB neutralization by the TcdB-binding-protein cocktail indicated that the optimal ratio at high total TcdB-binding protein concentrations was ≪1:0.8:1, primarily because pp1005 activity had a nonlinear (complex) relationship with concentration (see Supplemental Materials). However, at low total concentrations, pp1005 made a major contribution to cocktail potency and in vivo data showed that it was the most protease-sensitive component of the cocktail. Therefore, pp1005 content was increased to ensure potent activity of the cocktail over a broad concentration range, especially after exposure to intestinal proteases. The molar concentration of pp1005:pp1006:pp1007 in the final drug product was 0.1:0.5:1, which was associated with high potency over a wide concentration range.

### Phase 2 clinical efficacy of LMN-201 (RePreve Trial Part A Interim Results)

The RePreve Trial (study CDI02; NCT05330182) was designed to evaluate the efficacy of LMN-201 in preventing recurrent CDI. Here we report interim results of Part A of the RePreve Trial: a Phase 2 open-label assessment of LMN-201 with primary endpoints of safety and tolerability. This included pre-planned analyses, in coordination with the independent data and safety monitoring board, of primary cure and recurrence in the 21 participants who completed the Part A treatment regimen. Data cutoff for this analysis was May 21, 2025.

Participants received LMN-201 twice daily in conjunction with SoC antibiotics for 7 days, then were closely monitored for CDI recurrence for four weeks. We evaluated efficacy in relation to historical SoC controls in the MODIFY I/II trials^18^ for bezlotoxumab (NCT01241552 and NCT01513239), which had broadly similar eligibility criteria (Supplemental Table 5), SoC treatments, and endpoint definitions (while recognizing differences in era, sites, and supportive care).

Twenty-two Part A participants were enrolled between August 29, 2024, and February 14, 2025, and received one or more doses of LMN-201. One participant was lost to follow-up with no further contact. Twenty-one Part A participants completed 7 days of LMN-201 in addition to SoC antibiotics. Of these 21 participants, 16 (76%) had treatment-emergent adverse events within 30 days of the first dose of LMN-201, none of which were considered study drug-related. Most were mild to moderate gastrointestinal disorders and quickly resolved. Three participants (13.6%, 3/22) had serious adverse events that began after completion of 7 days of LMN-201 but within 30 days of the first dose of LMN-201. These resolved and were not considered study drug-related.

All 21 participants achieved initial clinical cure (21/21; 95% confidence interval (CI): 85–100%), higher than the initial clinical cure rates in the MODIFY I/II trials for both SoC antibiotics alone (80.3%; 621/773; 95% CI: 77–83%) and bezlotoxumab combined with SoC antibiotics (80.0%; 625/781; 95% CI: 77–83%) (p<0.001 for LMN-201 against SoC and SoC plus bezlotoxumab by naive two-sample test of proportions without multiplicity correction) (Figure 6)^18^. For Part A of the RePreve Trial, the sustained cure rate was defined as treatment success through four weeks with no new episode of diarrhea with a positive TcdB immunoassay occurring during the four-week observation phase; this captures the majority of recurrence events and aligns with the period most commonly used in past antibiotics trials. The sustained cure rate in RePreve Trial Part A was 90.5% (19/21; 95% CI: 71–97%), again higher than published rates for SoC antibiotics alone (59.5%; 460/773; 95% CI: 56–63%) and SoC antibiotics plus bezlotoxumab (68.9%; 538/781; 95% CI: 66–72%) (p<0.001 for LMN-201 against SoC and SoC plus bezlotoxumab by naive two-sample test of proportions without multiplicity correction).

**Figure 6:**
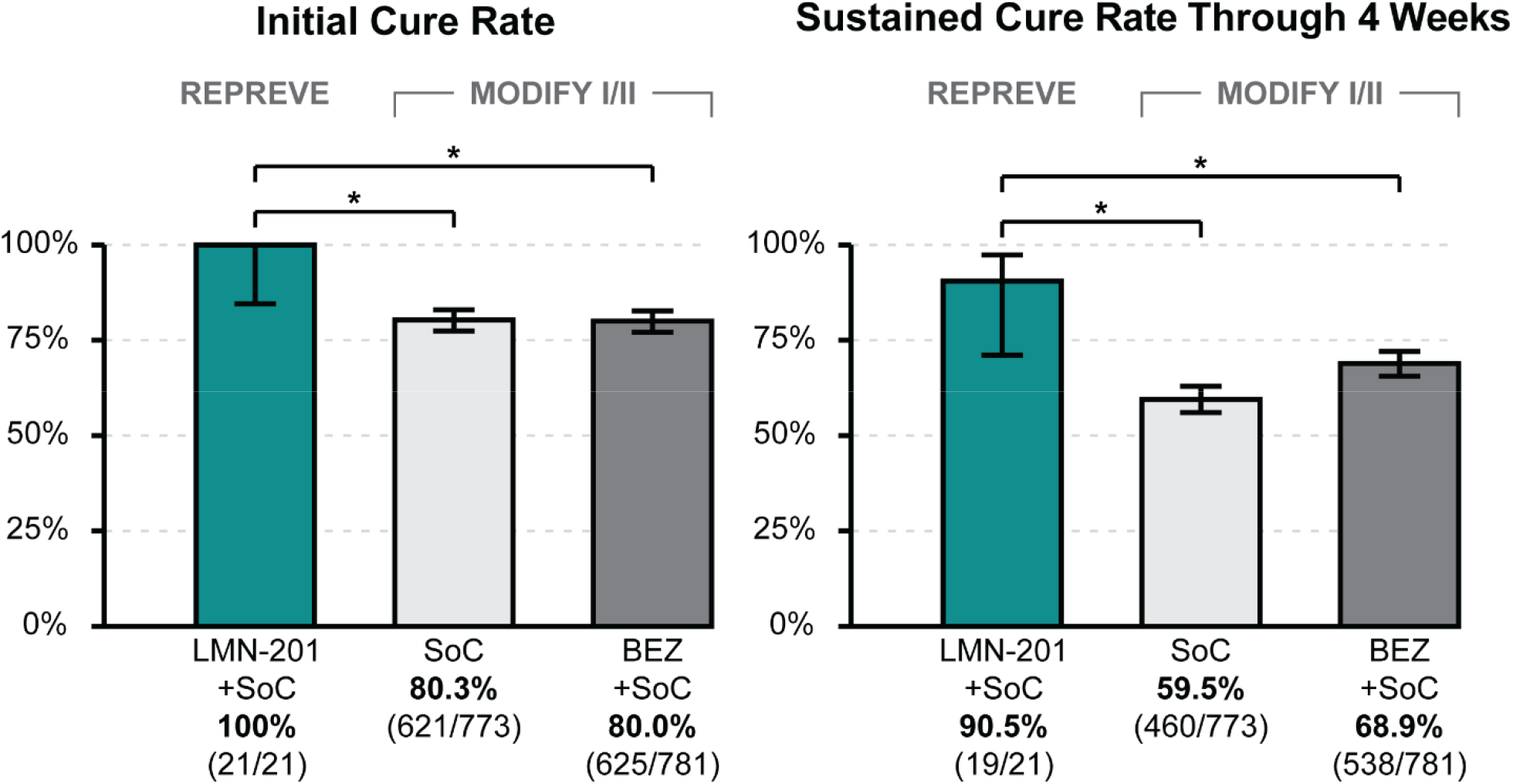
Phase 2 Clinical Efficacy of LMN-201 (RePreve Trial Part A). Initial clinical cure (left panel) and sustained cure through four weeks (right panel) comparing LMN-201+SoC, SoC alone and bezlotoxumab **(BEZ)**+SoC. Bezlotoxumab provided no additional benefit to initial clinical cure (left panel) above SoC alone and provided a modest improvement in sustained clinical cure through four weeks above SoC alone. In contrast, LMN-201 significantly improved both initial clinical cure rate and sustained clinical cure rate relative to SoC alone (historical data comparison). Error bars indicate 95% confidence intervals (Wilson method).

Although caveats still apply, cross-trial comparison with MODIFY I/II is appropriate because the RePreve study was intentionally designed to closely mirror those trials in key dimensions, including patient population, endpoint definitions, and use of standard-of-care antibiotics. Both programs evaluate the same clinical construct (administration of a targeted biologic therapy concomitantly with SoC antibiotics) and enroll adults with confirmed CDI receiving similar antibiotic regimens. Importantly, demographic characteristics and disease features are broadly comparable. While differences in practice patterns exist, these are well characterized, and overall concordance supports meaningful contextual comparison (Supplemental Table 5).

## Discussion

Many prevalent infectious, metabolic, and inflammatory disease processes that originate in the GI tract remain unaddressed or addressed only indirectly by systemic administration of biologic drugs. Orally delivering a therapeutic amount of a protein drug directly to its site of action in the GI tract is exceptionally challenging. The pharmacokinetic barriers to achieving this goal are primarily the high amounts of proteases in the GI tract that can quickly reduce the amount of a protein drug product below a minimally effective level, and the rapid rate of transit of ingested drugs through the intestine. Both can drastically decrease the magnitude and duration of therapeutic effect.

Here we showed that advantageous features of the spirulina biomanufacturing and drug delivery platform allowed us to achieve this goal: dose size, dose frequency, and potency together elevated to therapeutic levels the amount of drug product delivered to GI targets. High dose levels were possible by constitutively expressing, in spirulina, soluble, bioactive proteins in amounts ranging from approximately 0.5% to 3.4% of total biomass. Even at the low end, a palatable one-gram oral dose of dry spirulina biomass was sufficient to initially deliver a low micromolar concentration of drug product to the free-fluid compartment of the intestine (approximately 100 mL), which was orders of magnitude greater than the EC typically reported for cytokines, hormones, or antibodies^38,39^. Frequent dosing—which addressed the intrinsically transitory nature of GI drug delivery—was facilitated by spirulina’s simple, scalable GMP production environment.

A third advantage of the platform was the ease with which cocktail therapeutics were developed. The FDA requires assurance of safety and GMP manufacturing controls for biologic drugs prior to human testing. The relative simplicity of spirulina engineering and production processes—and the safety of oral delivery relative to injected—meant that even high-valence biologic cocktails could be developed on the platform for a fraction of the logistical intensity and time required for a single, conventional (injected) biologic drug molecule.

The high potency of the LMN-201 VHH cocktail is a form of Loewe synergy^40^ (also known as Bliss independence, which is a statistical enhancement of potency due to the independent binding probabilities of the individual VHHs)^41,42^. For the specific case of antibody cocktail therapeutics, we term this phenomenon “synthetic avidity”, wherein the simultaneous targeting of independent, non-overlapping epitopes by a cocktail of protein binders results in a multiplicative increase in neutralization potency—achieving values that mirror the high-avidity interactions seen in multimeric natural antibodies. Synthetic avidity is distinguished from classical avidity because it achieves these apparently high-affinity interactions through independent binding rather than physical linkage. This can be quantitatively modeled by studying the effects of VHH mixtures across a large range of TcdB concentrations (Supplemental Materials and Supplemental Figures 1, 2, and 3).

To our knowledge, LMN-201 represents a first-in-class human biologic rationally and intentionally engineered to utilize Bliss-independent multiplicative potency as a primary design principle. Unlike antibody cocktails that were designed for mutational breadth^43^, LMN-201 was developed to achieve synthetic avidity, reaching femtomolar neutralization thresholds through the coordinated effect of independent, non-overlapping binding proteins. The 200-fold decrease in the concentration of the biologic cocktail required to neutralize TcdB compared to any one of the individual proteins was essential for achieving sustained therapeutic levels at the distal intestine site of infection and likely contributed to the high response rates observed in the Phase 2 study.

The multiplicative gains in potency achieved by combining drugs that independently neutralize a target increase nonlinearly with drug concentrations and can be enormous when concentrations of each component substantially exceed their threshold of activity. Therefore, synergistically acting antibodies can be particularly useful for treating a disease in which the abundance of the causative agent is far above its minimal pathogenic level. For example, the concentration of C. difficile toxin B in patients typically exceeds its IC_50_ for epithelial toxicity by orders of magnitude. Hence, even highly potent single antibodies, with mid-picomolar EC_50_s, would need to be delivered at unrealistically high concentrations to reduce toxin activity below clinically significant levels. In contrast, drug cocktails, with apparently low femtomolar combinatorial EC_50_s, achieve therapeutic concentrations even when substantial amounts of the drug product are lost to proteolytic degradation. That the cocktail also offers broader strain coverage and protection against escape variants are additional advantages. Beyond infectious diseases, Bliss independence may also be useful for treating diseases with complex etiologies where single-agent interventions have only been moderately successful, for example inflammatory diseases of the gastrointestinal tract and cardiometabolic disease.

The fourth LMN-201 component—a bacteriophage-derived endolysin—not only broadens C. difficile strain coverage but also further enhances potency. While TcdB is the key virulence factor in CDI, approximately 5% of patients are infected by C. difficile strains that secrete TcdA but not TcdB^32^. Other virulence factors are also associated with CDI, including the C. difficile binary toxin (CDT), which is associated with increased 30-day mortality^44^. Inclusion of the lysin may offer protection against these non-canonical C. difficile strains, as well as novel strains with new toxin variants that might emerge in the future.

The lysin may also increase potency through synergy with vancomycin (the antibiotic most often prescribed for CDI); the two act in concert to disrupt C. difficile cell wall structure^28^. Also, these agents independently intervene at two steps in the pathway that controls virulence—C. difficile proliferation and toxin action on the gastrointestinal epithelium. The lysin thus complements the TcdB-binding proteins with a potential for orthogonal synergy to enhance LMN-201’s clinical real-world effectiveness.

The cocktail therapeutic described here also has important benefits for combating anti-microbial resistance, a problem that can emerge from—and hence place limitations on—the widespread use of potent anti-pathogen therapeutics, especially small-molecule antibiotics. The TcdB-binding proteins in LMN-201 recognize targets that are specific to the selected pathogen and therefore do not provoke emergence of resistance mechanisms in bystander microbes, including the normal intestinal flora, which often serves as a reservoir of antibiotic resistance genes. Our use of three VHHs that are all neutralizing on their own and bind to different sites on the TcdB toxin may decrease the likelihood of escape variants arising during therapy. Moreover, lysin escape mutations have never been observed experimentally^45,46^.

In addition to potency, dose size, and dose frequency, a fourth parameter affecting the oral delivery of protein drugs is stability, which in this case is primarily sensitivity to degradation by intestinal proteases. We observed a correlation between the sensitivity of a drug substance to major GI proteases and the amount recovered in the distal small intestine following oral administration. While we have not yet engineered the proteins in LMN-201 specifically to improve their protease resistance, this will be an important new direction for future drug development.

## Methods

### Development and characterization of LMN-201

LMN-201 is an investigational four-component protein cocktail comprising one C. difficile–specific endolysin and three TcdB-binding proteins derived in part from VHHs 5D, E3, and 7F previously reported. The endolysin component was derived from PlyCD. Lysins are bacteriophage-encoded enzymes which catalyze the cleavage of specific microbial cell wall components^28,47,48^. The VHHs were previously shown to bind and inactivate TcdB^9,29–31^. Binding of VHH 5D to TcdB prevents its pH-driven conformational change that enables pore formation and transit into the cytoplasm from internalized endosomes. VHH E3 binds to the glucosyltransferase domain of TcdB, interfering with the autoproteolysis step necessary for enzyme activation. VHH 7F also binds to the glucosyltransferase domain, inhibiting catalytic glucosylation/inactivation of Rho-family GTPases by the mature N-terminal glucosyltransferase.

The primary structures of each monomer subunit were—from amino to carboxy terminus—the maltose-binding protein chaperone (366 amino acids)^49^, a Gly-Gly-Gly-Gly linker, a 5HVZ dimerization domain (49 amino acids), a Gly-Ser-Gly-Ser-Gly-Gly linker, a TcdB-binding VHH (111–127 amino acids), a Gly-Ser-Gly linker and a C-terminal hexahistidine tag (total length, 543–562 amino acids) (Figure 1A). The 5HVZ dimerization domain was derived from the mammalian cAMP-dependent protein kinase A holoenzyme^50,51^.

Each final spirulina expression strain (SP1287, SP1308, SP1312 and SP1313) contained the corresponding gene of interest integrated into the spirulina chromosome at a single, pre-selected location, with no additional exogenous genetic material. Clonal spirulina filaments were isolated by micromanipulation, confirmed to be homozygous for the transgene, and expanded to generate frozen strain banks.

Expression levels of all four therapeutic proteins in spirulina were quantified using a capillary electrophoresis-based immunoassay. Binding kinetics were measured by biolayer interferometry using recombinant TcdB subdomains corresponding to the specific antigen-binding sites of each VHH.

### Toxin neutralization assay

Disruption of the actin cytoskeleton by TcdB causes changes in cell shape and cell attachment. In vitro, these cytoskeletal changes can be measured in real-time as a change in impedance using cells cultured on a surface embedded with microelectrodes^33,52^.

TcdB1, the toxin subtype most frequently present in worldwide C. difficile isolates from infected patients^53^, was used for primary analyses. TcdB1 was purified from C. difficile ribotype (RT) 087 (type strain 10463). Toxin concentrations tested spanned the range reported in patients with CDI^32^.

Genomic sequencing defined 12 distinct TcdB groups that differed from each other by 3% or greater^54^. 96% of the TcdB genes present in that dataset were defined as belonging to groups 1, 2, 3, and 5. Group 1 is the most clinically prevalent group worldwide, and group 2 is expressed by C. difficile ribotype 027, the single most prevalent strain type in the United States, and second most prevalent in Europe^5,53,55,56^.

### Engineering of spirulina

Spirulina strains without exogenous selection markers were generated as previously described^27^. Briefly, the endogenous spirulina gene WP_006618409.1, which confers kanamycin resistance, was replaced with a streptomycin resistance cassette to generate strain SP205. SP205 was subsequently transformed with constructs containing the gene of interest (anti-TcdB VHHs 5D, E3, and 7F; anti-TcdA VHHs AC1 and AH3; PlyCD or PlyCD_1–174_), WP_006618409.1, and flanking homology arms. Transformants were selected using kanamycin.

Single spirulina filaments were isolated and expanded to generate cell banks. Genomic DNA extracted from cells grown from the cell banks was analyzed by PCR, Sanger sequencing, and Illumina next-generation sequencing. Complete chromosomal segregation was confirmed, and sequences of the integrated gene of interest and WP_006618409.1 locus were verified.

### Next-generation sequencing of transformed spirulina

Genomic DNA was extracted from spirulina using a Blood and Tissue Kit (Qiagen) with minor modifications. Cells were resuspended in buffer ATL and digested with proteinase K (2 mg/mL) overnight at 56°C, followed by RNase A treatment (10 mg/mL, 20 minutes at 56°C). Buffer AL (400 µL) was added, and samples were centrifuged for 10 minutes at 13,000 × g. Supernatants were mixed with 250 µL of 100% ethanol and applied to DNeasy columns, which were washed once with 500 µL buffer AW1 and once with 500 µL AW2. DNA was eluted in molecular-grade water.

Purified genomic DNA was quantified using Qubit dsDNA High Sensitivity and Broad Range Kits (Thermo Fisher Scientific) and prepared for sequencing using the Illumina DNA prep kit. Sequencing was performed on a MiSeq instrument using a MiSeq Reagent Kit v2 (Illumina).

### Analysis for copy number determination

Transgene copy number in strains SP1287, SP1308, SP1312, and SP1313 was determined by comparing read depth of the integrated gene to 10 randomly selected normalization genes (ppaX, rpsA, chlM, degU2, radA, rbcL, sdhA, psbC, cysA1, smc6). A copy number of 1 ± 0.15 was used as the benchmark for complete single-copy genomic integration. As a control, a second set of ten randomly selected genes (thrB1, hisC1, isfD2, pilA, purB, apcE, nblA, gpmB, bicA2, cpcBA) was analyzed pairwise against the normalization gene set.

Read depth per base was calculated from Illumina data using samtools mpileup. (http://www.htslib.org/doc/). Median read depth across the transgene was normalized to the median read depth of the normalization genes to determine copy number per genome.

### Expression analysis of recombinant proteins in spirulina

Dried or wet spirulina biomass was resuspended in phosphate-buffered saline (PBS) containing EDTA-free protease inhibitor cocktail and 1 mM phenylmethylsulfonyl fluoride (PMSF; Thermo Fisher Scientific) at 2–10 mg/mL and lysed with 0.1 mm silica beads using a Precellys Evolution homogenizer (Bertin Technologies). Soluble extracts were clarified by centrifugation at 15,000 g for 30 minutes at 4°C.

Soluble recombinant protein levels were quantified using a capillary electrophoresis–based immunoassay (Jess, ProteinSimple) according to the manufacturer’s instructions. Extracts were diluted to 0.1–1 mg/mL biomass, and standard curves were generated using purified protein. Proteins were separated using a 12–230-kDa separation module and detected with a mouse anti-His-tag antibody (1:100; GenScript) followed by an anti-mouse IgG NIR secondary antibody (ProteinSimple). Data analysis was performed using the ProteinSimple Compass software and Microsoft Excel.

### Protein purification from spirulina biomass

The TcdB-binding proteins (pp1005, pp1006, and pp1007) and lysin PlyCD_1–174_ (pp1092) were purified from spray-dried spirulina biomass using a two-step Ni–NTA affinity and size-exclusion chromatography workflow. Biomass was resuspended in PBS with protease inhibitors (30 mL/g) and lysed by high-pressure homogenization (Microfluidics LM20, 12,000 psi). Lysates were clarified by centrifugation at 18,500 × g for 30 minutes at 4°C and sequentially filtered through coffee filter and 0.22 µm membrane filters.

Clarified lysates were applied to 5 mL HisTrap columns (Cytiva) equilibrated in 50 mM Tris (pH 8.0), 300 mM NaCl and 20 mM imidazole (Thermo Fisher Scientific). Eluted proteins were further purified by size-exclusion chromatography using a HiLoad 16/600 Superdex 200 pg column on an ÄKTA Pure system. Protein purity and size were assessed by analytical size-exclusion chromatography (Superdex 200 Increase 10/300) and SDS–PAGE under reducing and non-reducing conditions. Purified proteins were >90% pure and exhibited expected molecular weight distributions.

### Cell lines and toxins

Vero cells (ATCC) were maintained in Dulbecco’s Modified Eagle Medium supplemented with 10% heat-inactivated fetal bovine serum and 1% penicillin–streptomycin (Gibco) at 37°C and 5% CO₂. C. difficile toxin TcdB-10463 was expressed in Bacillus megaterium and purified by Ni-NTA affinity and ion-exchange chromatography. Purified TcdB-027, TcdB-1470, and TcdB-078 were obtained from tgcBIOMICS. Bezlotoxumab was obtained from Henry Schein Medical.

### Expression of recombinant TcdB toxin, inactive TcdB, and TcdB subdomains in bacterial systems

Active and inactive TcdB were expressed in Bacillus megaterium and purified as previously described, with minor modifications^57^. Transformant strains were provided by the laboratory of C. Shoemaker (Tufts University). Bacteria were cultured in LB medium containing tetracycline (10 µg/mL) and expanded in Super Broth supplemented with tetracycline. Protein expression was induced with 0.5% (w/v) xylose at OD₆₀₀ 0.3–0.4, and cultures were harvested after 16 hours.

Cell pellets were resuspended in lysis buffer (50 mM sodium phosphate, 300 mM NaCl, 20 mM imidazole, 0.5 mM EDTA, pH 8.0) and lysed by sonication. Clarified lysates were purified by Ni–NTA affinity chromatography using 5 mL HisTrap columns on an ÄKTA Pure system. Bound proteins were eluted with an imidazole gradient, and fractions were analyzed by SDS–PAGE. Selected fractions were pooled, concentrated, further purified by ion-exchange chromatography (HiTrap Q XL), buffer-exchanged into PBS, and assessed by SDS–PAGE under reducing and non-reducing conditions. Protein concentration was determined by absorbance.

TcdB subdomains for VHH binding^30^ were designed for bacterial expression with an N-terminal hexahistidine tag, maltose-binding protein, and TEV protease cleavage site, and a C-terminal Avi-tag for enzymatic biotinylation. Based on the structural information, we designed the 5D-binding region and the delivery and receptor-binding domain for expression using residues 1092-1433. VHHs 7F and E3 bind regions on the glucosyltransferase domain. A fragment containing this region, from amino acid positions 2 through 543 of TcdB, was subcloned into a modified pET28 b (+) vector where the kanamycin bacterial resistance gene was replaced with ampicillin bacterial resistance gene. Sequence-verified plasmids were transformed into E. coli BL21(DE3) (New England Biolabs).

Protein was expressed using autoinduction protocol^58^. Cultures were grown at 37°C followed by 16°C, harvested by centrifugation, and stored at −20°C. Pellets were resuspended in buffer containing 50 mM Tris pH 8.0, 300 mM NaCl, protease inhibitors and 1 mM PMSF. Cells were lysed by high-pressure homogenization. Clarified lysates were purified by Ni–NTA affinity chromatography, followed by size-exclusion chromatography on a Superdex 200 column equilibrated in PBS. Purity was assessed by SDS– PAGE. Purified proteins were biotinylated using BirA ligase according to the manufacturer’s instructions (Avidity, Inc.).

### Assessment of binding to TcdB subdomains with biolayer interferometry

Biolayer interferometry measurements were performed on the Octet RED96 system (Sartorius) using High-Precision Streptavidin biosensors (Sartorius). Biosensors were hydrated with PBS, pH 7.4 at room temperature for 10 minutes on microplate 96-well flat-bottom (Greiner). All kinetics experiments were performed at 30°C with 500 rpm agitation in the kinetics module. Biosensors were dipped into PBS containing wells for 60 seconds, followed by loading with enzymatically biotinylated TcdB antigen fragments at 10 µg/mL for 200 seconds to achieve ∼0.6–1 nm response. Loading was quenched by incubating biosensors in 50 µM biocytin (Sigma Aldrich) for 60 seconds. Baselines were established by incubating antigen-loaded biosensors in kinetics buffer (PBS + 0.02 % Tween 20, 0.1 % BSA, 0.05 % sodium azide) for 300 seconds. The rate of association was measured by incubating antigen-loaded biosensor tips into three-fold dilution series of monomeric (pp1239, pp1238, and pp1240, respectively) or dimerized (pp1005, pp1006, and pp1007, respectively) 5D, E3, and 7F VHHs. The concentration series used to determine binding kinetics of pp1238 were 93.6, 31.2, 10.4, 3.47, 1.16, and 0.36 nM. Concentrations used to determine binding kinetics of pp1239 were 93.6, 31.2, 10.4, 3.47, and 0.38 nM. Concentrations used for pp1240 were 92.2, 30.7, 10.2, 3.41, 1.14, and 0.38 nM. pp1005 was assayed at 16.2, 5.39, 1.8, 0.60, 0.20, and 0.07 nM. Concentrations used for pp1006 were 16.8, 5.6, 1.87, 0.62, 0.21, and 0.07 nM. pp1007 was assayed at concentrations of 84.9, 28.3, 9.43, 3.14, 1.05, and 0.35 nM. Analyte-bound biosensors were dipped into kinetics buffer for 300–600 seconds to measure rate of dissociation. Kinetic analysis was performed using the HT 11.1.1.39 Data Analysis module (Sartorius). Results were double referenced. Both association and dissociation steps were used in 1:1 binding global data fitting model.

### Toxin-neutralization assay

Toxin-neutralization assays for TcdB1.1, TcdB2.1, TcdB3.1, and TcdB5.1 were performed using the xCELLigence Real-Time Cell Analysis MP system (Agilent). Cell health was measured in real-time using an impedance-based readout, in which changes in electrical impedance generated by adherent cells on gold microelectrodes in 96-well E-plates are reported as a unitless cell index. TcdB-induced cytotoxicity was quantified as a reduction in cellular adhesion reflected by decreased Cell-index values.

Vero cells were seeded at 1 × 10⁴ cells per well in 100 μL DMEM supplemented with 10% fetal bovine serum and 1% penicillin–streptomycin in Agilent 96-well E-plates. Plates were equilibrated for 30 minutes at room temperature and transferred to the xCELLigence MP instrument maintained at 37°C with 5% CO₂ (Caron). Cell-index measurements were acquired at 15-minute intervals for 20–24 hours.

TcdB toxins were serially diluted and pre-incubated at 37°C for 1 hour with purified TcdB-binding protein, lysin, or bezlotoxumab. The assay was paused, culture medium was removed, and 100 μL of toxin–protein mixtures was added to each well. Cell-index monitoring was resumed and continued for at least 24 hours.

Cell-index values at 24 hours post-treatment were normalized to the final measurement obtained immediately prior to treatment addition. The resulting normalized cell index was plotted as an adhesion score using GraphPad Prism, and IC₅₀ values of TcdB were determined by nonlinear regression analysis.

### TcdB’s effect on cell adhesion and inhibition of TcdB by TcdB-binding proteins

Toxin-neutralization assay data were imported, normalized, and analyzed in R (version 4.1.1; R Foundation for Statistical Computing). Nonlinear regression analyses were performed using the minpack.lm routine in the nlsLM package, which applies the Levenberg-Marquardt algorithm to perform bounded minimization^59^.

### Lysin activity assay using Bacillus subtilis SL4

The lytic activity of pp1092 against Bacillus subtilis SL4 was assessed using a turbidity-reduction assay adapted from previously described methods to a 96-well plate format^28^. B. subtilis SL4 cultures grown to mid-log phase in brain-heart infusion medium (37°C, 250 rpm) were harvested and washed twice with PBS by centrifugation (6,000 × g, 10 minutes, room temperature). Bacterial suspensions were adjusted to an OD₆₀₀ of 1.0 and incubated with 1 mg of digested or undigested pp1092 in a final volume of 200 μL at 37°C with shaking.

Lytic activity was quantified by measuring the reduction in turbidity (OD₆₀₀) over time. Each test sample was analyzed alongside five, three-fold serial dilutions of pp1092 standards and fit using a modified five-parameter nonlinear logistic model. The time to 50% reduction in OD₆₀₀ (TC₅₀) was estimated using a four-parameter nonlinear logistic model with lysin concentration as the independent variable. Standard curves were generated by linear regression analysis in Microsoft Excel using TC_50_ values plotted against known pp1092 concentrations. The activity of unknown samples was determined by interpolating their TC_50_ values from the standard curve and was expressed as the equivalent concentration of pp1092.

### Lysin activity assay using C. difficile

C. difficile strain ATCC 43255 was grown to mid-log phase (OD_600nm_ = 0.5) at 37°C under anaerobic conditions in reduced brain heart infusion medium (BD Biosciences) supplemented with 0.5% (wt/vol) yeast extract and 0.1% (wt/vol) L-cysteine. Cultures were harvested by centrifugation (4,000 rpm, 15 minutes), washed, and resuspended to an OD_600_ of 2.0 in 50 mM sodium phosphate buffer (pH 7.0).

Lysin activity was assessed using a turbidity-reduction assay in 96-well flat-bottom, untreated polystyrene microtiter plates (Corning). Wells were preloaded with 100 μL of buffer containing lysin at final concentrations ranging from 4 to 256 μg/mL or buffer alone (untreated control). Bacterial suspensions (100 μL) were added to each well, and OD₆₀₀ was measured at 60-second intervals for 15 minutes at 37°C using a SpectraMax M5 microplate reader (Molecular Devices).

### Lysin protease digestion and quantification

Lysin proteins (0.2 μg/μL) were incubated in digestion buffer (20 mM Tris, 150 mM NaCl, 3 mM CaCl₂, pH 6.4) with trypsin (Promega), chymotrypsin (MP Biomedicals), or elastase (MP Biomedicals) at final concentrations of 0.01 or 0.1 mg/mL. Reactions were incubated at 37°C with shaking at 200 rpm for 1 hour and immediately quenched by addition of 1 mM PMSF and 1× Pierce Protease Inhibitor Mini Tablets, EDTA-free (Thermo Fisher Scientific). For elastase-treated samples, 1 mM elastase inhibitor III (MilliporeSigma) was also added.

Protease digestion was visualized by resolving 2 μg of digested and undigested protein on gradient BOLT-PAGE gels (Thermo Fisher Scientific) run at 200 V for 20 minutes and stained with Coomassie Blue. Band intensities were quantified using ImageJ, with background subtraction performed using identically sized regions of interest and signal normalized to the corresponding undigested protein band for each condition.

### TcdB-binding protein protease digestion for ELISA

In vitro protease digestion of TcdB-binding proteins was performed in a matrix of soluble lysate from wild-type spirulina (SP003) prepared in Bis-Tris buffer (pH 6.0). Digestion reactions were treated with trypsin, chymotrypsin, or pancreatin (Sigma Aldrich). TcdB-binding proteins in PBS were diluted into SP003 lysate, Bis-Tris buffer, and protease to yield final concentrations of TcdB-binding protein, 20 mg/mL SP003 lysate, and protease corresponding to “fed” or “fasted” conditions. This concentration of SP003 lysate and TcdB-binding protein approximates a 1% (dry weight) expression level in spirulina.

For trypsin and chymotrypsin digestions, proteases were used at 0.1 mg/mL (“fed”) or 0.01 mg/mL (“fasted”) concentrations. Pancreatin was used at 0.1 mg/mL. Reactions were incubated at 37°C for 3 hours and quenched by addition of an equal volume of stop buffer containing 2 mM PMSF and 2× Pierce Protease Inhibitor Cocktail in PBS. Following protease neutralization, TcdB-binding proteins were present at a final concentration of 0.1 mg/mL. Samples were kept on ice until ELISA analysis.

### TcdB-binding protein quantitation by ELISA

Binding activity of digested TcdB-binding proteins was quantified by ELISA. Nunc MaxiSorp plates (Thermo Fisher Scientific) were coated with inactive TcdB (2 μg/mL; 100 μL per well) in carbonate– bicarbonate binding buffer and incubated overnight at 4°C. Plates were washed three times with PBS containing 0.05% Tween-20 (PBS-T) and blocked for 1 hour with 300 μL per well of blocking buffer (PBS-T supplemented with 5% BLOTTO).

Undigested and protease-digested TcdB-binding proteins were serially diluted in blocking buffer using low-binding 96-well plates. Six dilutions of undigested TcdB-binding protein were used to generate standard curves, and two dilutions of each digested sample were analyzed for quantification by interpolation. Samples (100 μL per well) were added to coated plates and incubated for 1 hour at room temperature with shaking. Plates were washed as described above, followed by incubation with horseradish peroxidase (HRP)-conjugated anti-camelid VHH antibody cocktail (GenScript) diluted 1:50,000 in blocking buffer (100 μL per well) for 50 minutes with shaking.

After washing twice with PBS-T and once with PBS, 100 μL per well of TMB Ultra substrate (Thermo Fisher Scientific) was added and plates were developed in the dark at room temperature for 10 minutes with shaking. Reactions were quenched with 50 μL of 2.5 M sulfuric acid, and absorbance was measured at 450 nm using a SpectraMax M5 microplate reader (Molecular Devices). All digestion conditions were analyzed in duplicate across three independent experiments.

### Calculation of TcdB-binding protein protease resistance

TcdB-binding protein concentrations in protease-digested samples were determined by interpolation from standard curves generated using undigested TcdB-binding protein. Protease resistance was expressed as the percentage of remaining TcdB-binding protein activity, calculated by normalizing the interpolated concentration of each digested sample to the corresponding undigested control. Calculations were performed for each replicate independently, and data are reported as mean ± s.d. across replicates.

### Preparation of C. difficile spores for mouse studies

C. difficile strain M7404 (a ribotype 027 clinical isolate from Canada) was cultured on HIS-T agar (3.7 % (w/v) heart infusion (HI) (Oxoid), 0.5 % (w/v) yeast extract, 1.5 % (w/v) agar supplemented with 0.375 % (w/v) glucose, 0.1 % (w/v) L-cysteine, and 0.1 % (w/v) sodium taurocholate (New Zealand Pharmaceuticals)) under anaerobic conditions (10% H₂, 10% CO₂, 80% N₂) at 37°C in an anaerobic chamber (Don Whitley Scientific).

For spore preparation, cultures were subcultured on HIS-T agar and inoculated into 500 mL tryptone– yeast broth (3.0% tryptone, 2.0% yeast extract, and 0.1% sodium thioglycolate), followed by anaerobic incubation at 37°C for 10 days. Spores were harvested by centrifugation (10,000 g, 20 minutes, 4°C), washed five times with chilled distilled water, and heat-shocked at 65°C for 20 minutes. Spore titers were determined by plating on HIS-T agar. Spores were diluted to 1 × 10⁶ CFU/mL in PBS containing 0.05% Tween-80 for mouse infection studies.

### Mouse study design

Male C57BL/6J mice were infected with C. difficile spores 24 hours after oral gavage with one of the following treatments: vancomycin (positive control), wild-type spirulina (negative control), Cocktail M1, or Cocktail M2. Cocktail M1 comprised equal proportions of spirulina strains SP1308, SP1312, and SP1313 (expressing pp1005, pp1006, and pp1007, respectively) and SP1286 (expressing full-length lysin PlyCD, pp1093; Supplemental Figure 4). Cocktail M2 was identical to Cocktail M1 but omitted SP1286. Treatments were administered once daily following initial C. difficile challenge on day 0.

All animal procedures were conducted in accordance with Victorian State Government regulations and were approved by the Monash University Animal Ethics Committee (AEC no. 21731). Male C57BL/6J mice (6–7 weeks old) were assigned to treatment, positive control, or negative control groups, and housed in groups of five.

To induce susceptibility to C. difficile, mice were pretreated with an antibiotic cocktail in drinking water containing kanamycin (0.4 mg/mL), gentamicin (0.035 mg/mL), colistin (850 U/mL), metronidazole (0.215 mg/mL), vancomycin (0.045 mg/mL), and cefaclor (0.3 mg/mL) for 7 days, followed by cefaclor alone (0.3 mg/mL) for an additional 4 days.

Spirulina formulations and controls were resuspended in PBS (5% wt/vol) by repeated mixing using an 18-gauge blunt needle and 1-mL syringe for 2.5 minutes and administered within 10 minutes of preparation. Vancomycin was diluted to 6.25 mg/mL before administration. Beginning 1 day prior to infection (day −1), mice received once daily oral gavage of 200 μL treatment for the first 2 days, followed by 100 μL once daily for the remainder of the study.

On the day of infection (day 0), antibiotic pretreatment was discontinued, mice were switched to plain water, and infection (1 × 10⁵ spores per animal) was initiated 3 hours after treatment dosing.

Mice were monitored daily for clinical signs of infection, including weight loss, and were euthanized upon reaching ≥15% body weight loss, predefined humane endpoints, or at study termination (day 4). As a surrogate measure of colonization, fecal samples collected at 24 hours post-infection and at euthanasia were analyzed to quantify C. difficile spore shedding. Outcomes in mice treated with spirulina therapeutic cocktails and the vancomycin positive control were compared with those in the wild-type spirulina control group.

### Quantification of spore shedding from mouse feces

Fecal pellets were collected at 24 hours post-infection and at euthanasia, weighed, and resuspended in PBS (100 mg/mL). Samples were heat-shocked at 65°C for 30 minutes and plated for spore enumeration on HIS agar supplemented with 0.1% (wt/vol) L-cysteine, 0.1% (wt/vol) sodium taurocholate, 0.375% (wt/vol) glucose, 250 μg/mL D-cycloserine, 8 μg/mL cefoxitin, and 10 μg/mL erythromycin.

Colonization data were analyzed by fitting linear models to log₁₀-transformed CFU/g values at each landmark time point, with treatment included as a potential explanatory factor. The coefficient for wild-type spirulina represents the mean log₁₀ CFU/g for that group, and coefficients for other treatments represent differences in mean log₁₀ CFU/g relative to wild-type spirulina. Standard errors, t-statistics, and two-tailed P values (α = 0.05) are reported.

### Analysis of clinical outcomes in the mouse study

Maximum weight loss was analyzed by fitting linear models with treatment included as a potential explanatory factor. The coefficient for wild-type spirulina represents the mean maximum weight loss in that group, and coefficients for other treatments represent differences in mean maximum weight loss relative to wild-type spirulina. Standard errors, t-statistics, and two-tailed P values (α = 0.05) are reported.

Analysis of survival data was performed on tabulated number at risk and number of events at each time. Data were compared for each treatment to wild-type spirulina results and a P value was computed using the log-rank statistic for each comparison.

### Hamster study challenge reagent preparation and confirmation

C. difficile strain R20291 (NAP1/BI/027) spore stock was heat-shocked at 65°C for 30 minutes and titered on TCCFA plates, yielding 5 × 10⁵ CFU/mL in PBS (5 mL total volume). The stock was diluted to 15,000 spores per 50 μL and further diluted 1:1 with sterile PBS containing 5% sucrose to achieve a final challenge dose of 15,000 spores in 100 μL per animal.

To confirm the administered dose, 100 μL of the final inoculum was serially diluted (ten-fold) and plated on TCCFA agar. Plates were incubated anaerobically at 37°C for 48 hours, colonies were enumerated, and the challenge dose was calculated based on CFU counts. The confirmed inoculum was 1.5 × 10⁴ CFU per 100 μL.

### Hamster study design

The Golden Syrian hamster model of CDI has been previously used to evaluate orally delivered antibodies targeting C. difficile toxins^60^. Experimental groups included Cohort 1 (wild-type spirulina control) and Cohort 2 (Cocktail H1), consisting of spray-dried spirulina biomass of strains expressing TcdB-binding proteins pp1005, pp1006, pp1007, two TcdA-binding proteins of similar construction (His-tagged VHHs expressed with maltose-binding protein on 5HVZ dimerization scaffolds)^29^, and pp1092 (PlyCD_1–174_). Anti-TcdA VHHs were included because TcdA contributes significantly to disease severity in hamsters, unlike in humans^8^.

Hamsters received wild-type spirulina or Cocktail H1 twice daily beginning 1 day prior to infection and continuing until humane endpoint or study completion (12 days post-infection). Animals were challenged by oral gavage with 1.5 × 10⁴ C. difficile R20291 spores.

Golden Syrian hamsters (Envigo) were examined upon arrival and acclimated for 2 days prior to study initiation. Animals underwent physical examination and were randomly assigned to treatment groups with adjustment to balance body weight. Hamsters were fed ad libitum with sterilized 2018S Teklad Global 18% Rodent Diet (Envigo) and sterile water.

To induce susceptibility to CDI, clindamycin (15 mg/mL in saline) was administered intraperitoneally at 30 mg/kg once daily, for three consecutive days. During antibiotic treatment, cage bottoms were changed daily to minimize recolonization via coprophagy.

### Quantification and analysis of cecal C. difficile colonization in hamsters

Cecal contents were collected at necropsy upon reaching humane endpoints or at study termination. C. difficile burden was quantified by quantitative RT-PCR of DNA extracted from cecal samples and interpolated against a standard curve generated from DNA isolated from vegetative C. difficile cells. Group comparisons were performed using Wilcoxon rank-sum (Mann–Whitney) test with continuity correction.

### LMN-201 sample preparation for potency quantitation

Spray-dried spirulina biomass (LMN-201 drug substance, LMN-201 drug product, or wild-type control, as applicable) was weighed into bead-beating tubes. Lysing Matrix B was added and briefly vortexed to distribute the beads within the dry powder. Ice-cold lysis buffer was then added, and samples were maintained on ice. Cells were lysed using a Precellys Evolution homogenizer (three cycles of 7,500 rpm for 30 seconds, with incubation on ice between cycles). Lysates were clarified by centrifugation at 12,000–18,000 × g for 10 minutes at 2–6°C, and supernatants were collected for analysis.

### LMN-201 lysin potency quantitation

Clarified lysates, thawed single-use internal control, and thawed single-use protein calibrator were maintained on ice prior to assay setup. Samples, wild-type spirulina lysate, and internal controls were prepared at 30 mg biomass/mL in lysis buffer. Wild-type lysate was used as the assay matrix for preparation of standard curves and sample dilutions.

100 μL of each sample, internal control, and standards were added to a 96-well plate following the block-effect mitigation pattern. A flash-frozen mid-log B. subtilis substrate was thawed and diluted into 12 mL assay buffer stored at 25°C. Cell substrate (100 μL) was added to each well to initiate the reaction, yielding a final lysed biomass concentration of 75 µg/mL (1:400 dilution; SP1287b drug substance) or 300 µg/mL (1:100 dilution; LMN-201 blended drug product).

Plates were analyzed without lids using a NEPHLOstar nephelometer with kinetic reads for 180 minutes. Each cycle consisted of 10 seconds double-orbital shaking (200 rpm) followed by turbidity measurement at 635 nm at 70-second intervals. Raw result report files were used for potency analysis.

### LMN-201 TcdB-binding protein potency quantitation

Therapeutic TcdB-binding protein concentrations were quantified using MSD. TcdB fragments were engineered based on published VHH–TcdB co-crystal structures to generate three truncated antigens, each selectively binding one TcdB-binding protein. Mutant TcdB fragments were conjugated to MSD linkers and immobilized in multiplex format within 96-well MSD plates.

Following antigen immobilization and washing, reference standards and sample dilutions were added and incubated. Plates were washed and incubated with a tri-clonal primary antibody recognizing the TcdB-binding proteins, followed by a SULFO-TAG–conjugated secondary antibody. After final washes, read buffer was added and plates were analyzed by MSD. TcdB-binding protein concentrations were interpolated from reference standard curves.

Assay compatibility was evaluated for blended drug product. For pp1092, lysates from VHH-expressing strains were spiked with pp1092 and assessed for recovery; percent recovery was within 20% of expected values, indicating no interference (Supplemental Table 3). For pp1005, pp1006, and pp1007, serial dilutions were quantified individually and in the presence of the other TcdB-binding proteins at maximal test concentrations. Accuracy was defined as 80–120% recovery. Valid assay ranges were defined as 160–0.66 pM (pp1005), 400–1.64 pM (pp1006), and 400–4.10 pM (pp1007) (Supplemental Figure 5). Sample concentrations were required to fall within the validated assay range. This assay enabled independent quantitation of each TcdB-binding protein for potency determination.

### Ileostomy GI pharmacokinetics clinical trial

Study CDI01 (NCT04893239) was a Phase 1, multi-site, open-label, non-randomized exploratory study designed to assess the GI pharmacokinetics of LMN-201. Healthy individuals with stable ileostomies received LMN-201 formulated in enteric-coated capsules (Lonza Capsugel Size 00 Vcaps Enteric). Ileostomy output was collected for quantitation of LMN-201 therapeutic protein components.

All participants received one enteric-coated capsule containing 500 mg SP1287 biomass (expressing pp1092). Three additional capsules (500 mg each) containing spirulina strains expressing individual TcdB-binding proteins were administered as follows: SP1308 (n=2), SP1312 (n=2), SP1313 (n=2). Six participants received 500 mg of each TcdB-binding protein strain in addition to SP1287; however, samples from only two were available for testing.

Ostomy fluid was collected hourly, including pre-dose samples. TcdB-binding protein concentrations were quantified using a modified TcdB toxin-neutralization assay in Vero cells, with concentrations interpolated from standard curves prepared in control ostomy fluid. Concentrations of pp1092 were determined using a turbidity reduction assay with interpolation from standard curves made in control ostomy fluid.

The minimum therapeutic concentration (MTC) for each TcdB-binding protein was defined as the concentration of the LMN-201 cocktail needed to reduce levels of active TcdB by 100-fold. This was the decrease in TcdB burden achieved by SoC antibiotics^33^ and the fold difference in TcdB levels between patients acutely ill with CDI and patients without disease recurrence^32^. Note that this is a clinically relevant definition of efficacy, and not precisely identical to the MEC described above. The MTC for PlyCD was defined as the concentration of lysin required to lyse 1×10^11^ C. difficile bacteria (approximating the C. difficile bioburden in CDI) within 1 hour^34–37^.

Total soluble TcdB-binding protein recovery was calculated as the sum of TcdB-binding protein amounts measured across hourly samples and expressed as a percentage of the administered dose.

### Quantification of pp1092 in ostomy fluid

Concentrations of pp1092 (PlyCD_1–174_) in ostomy fluid were quantified using a turbidity reduction assay. Clarified supernatants of sample and control ostomy fluids were diluted 1:20 in PBS. Purified pp1092 was diluted to 16 µg/mL in as many control fluids as existed: T_arrival_, T_0_, and T_exit_ (final timepoint collected) and then diluted into the same ostomy dilution to make standard curves.

100 µL of diluted samples and standards were added to a 96-well plate, followed by 100 µL of flash-frozen B. subtilis cells (OD₆₀₀ = 2.0). Plates were incubated at 37°C, and turbidity (OD₆₀₀) was measured every 2 minutes for 3 hours.

Data were analyzed using GraphPad Prism 10. Turbidity-versus-time curves for standards and samples were fit using four-parameter logistic (4-PL) regression. Curves from positive samples (identified by green color) were overlaid with each standard to qualitatively identify the most parallel standard for interpolation. Chosen standard curve 4-PL fit IC_50_ values were plotted against PlyCD_1–174_ concentration using a 4-PL fit. Sample IC_50_ values were interpolated from the standard curve and corrected for dilution to determine PlyCD_1–174_ concentration.

### Quantification of TcdB-binding protein in ostomy fluid

Toxin-neutralization assays mentioned above, against TcdB1.1, were used to quantify the active TcdB-binding proteins in ostomy fluid collected in the intraluminal pharmacokinetic trial. For patients who received lysin and a single TcdB-binding protein, the corresponding purified TcdB-binding protein was serially diluted to be used as standards. For patients who received lysin and all three TcdB-binding proteins, three purified TcdB-binding proteins at their delivery ratio (1:3:10) were mixed and serially diluted to be used as standards. All the samples and standards were tested against 0.025 pM of TcdB. Measured adhesion score with each sample was interpolated to the standard curve to obtain the concentration of TcdB-binding protein. For patients 201-001 and 201-002, purified pp1005 at five concentrations was tested against serially diluted TcdB. A linear standard curve was generated with EC_50_of TcdB and concentrations of pp1005. Ostomy fluid samples were also diluted in matrix and tested against serially diluted TcdB. EC_50_ was interpolated to the standard curve to obtain the concentration of active pp1005 in the samples. To control for the matrix effect, fluid collected at the time of arrival for each patient was diluted 50-fold with cell culture medium and used to dilute samples and standards. The exception is patient 201-005, from whom the fluid at arrival was unavailable, therefore fluid from patient 201-006 at arrival was used to generate matrix.

### Phase 2 clinical trial—methods

The RePreve Trial (study CDI02; NCT05330182) was designed to evaluate the efficacy of LMN-201 in treating and preventing CDI. Trial endpoints and patient inclusion/exclusion criteria were aligned with the MODIFY I/II trials for bezlotoxumab (NCT01241552 and NCT01513239) to improve comparability (see Supplemental Table 5).

Part A of the RePreve Trial—the Phase 2 segment—was conducted at 13 sites across the U.S. from August 2024 to December 2025 (first patient first visit: August 29, 2024; last patient first visit: February 14, 2025; last patient last visit: August 20, 2025) in accordance with Good Clinical Practice guidelines and the provisions of the Declaration of Helsinki. Sample size for Part A was determined in consultation with the FDA; the primary endpoints for Part A were safety and tolerability. Primary cure and recurrence in coordination was also conducted in coordination with the data and safety monitoring board. The protocol and amendments were approved by a centralized institutional review board (Advarra, Columbia, MD, USA). Written informed consent was obtained from all participants. The RePreve Trial was designed by representatives of Lumen Bioscience and its academic advisors. All data were collected by investigators and associated site personnel, analyzed by statisticians at Lumen and its professional advisors, and interpreted by the authors.

Participants were adults with recently diagnosed CDI who were receiving or planned to receive standard-of-care oral antibiotic therapy (fidaxomicin, vancomycin, or metronidazole, selected by the treating physician) for 10–28 days. CDI was defined as a history of at least three loose or watery bowel movements (Bristol stool types 5 to 7) within 24 hours and a positive stool test for TcdB obtained within seven days before enrollment, with no alternative explanation for symptoms. Diagnostic testing was performed at local laboratories using validated toxin immunoassays. Stool samples were also collected for central laboratory analyses, including microbiologic and exploratory assessments. Detailed inclusion and exclusion criteria are provided in Supplemental Table 4.

LMN-201 was administered orally twice daily for 7 days, initiated as soon as feasible after the start of antibiotic treatment. Participants were followed through a four-week observation period and a subsequent follow-up phase for safety assessments. Participants recorded daily bowel movements, and new episodes of diarrhea were monitored through scheduled contacts and unscheduled visits. Safety assessments included monitoring of adverse events, laboratory testing, and collection of serious adverse events throughout the study period, in accordance with the protocol.

Twenty-three participants enrolled and 21 participants completed SoC antibiotics (metronidazole, vancomycin, or fidaxomicin) plus seven days of LMN-201 (Supplemental Figure 6). Analyses were conducted in all evaluable completed participants (those who received seven days of LMN-201 and had a confirmed diagnosis of CDI at baseline) with a data cutoff date of May 25, 2025. Efficacy was assessed descriptively in this treated population, which included participants receiving LMN-201 in combination with standard-of-care antibiotics, with outcomes compared to historical benchmarks. Data were taken from the unlocked electronic case report form (eCRF) database as of May 1, 2025. They were source-verified but remain subject to change until database lock at the trial’s conclusion. Safety and tolerability were assessed in all participants who received at least one dose of LMN-201, with no dose-related serious or severe adverse events observed. No discontinuations were considered drug-related.

## Supplemental Materials

### Derivation of Bliss-independence potency shifts for TcdB-binding protein mixtures

Derivation of the synthetic avidity model begins with the pharmacology model of TcdB1 action on cell adhesion described in the main text:

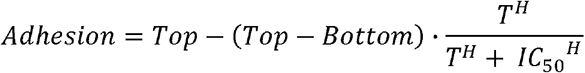

where Adhesion indicates the results of the toxin neutralization assay; **Top** and **Bottom** are the highest (no TcdB1) and lowest (maximal TcdB1) assay values; **T** is the concentration of TcdB1; **IC_50_** is the concentration of TcdB1 eliciting half maximal inhibition in Adhesion; **H** is the steepness of the concentration-response relationship.

In the context of one or more orthosteric inhibitors, only a fraction, **f_T_**, of T is available to bind and affect the target cells. The model for TcdB1 is then:

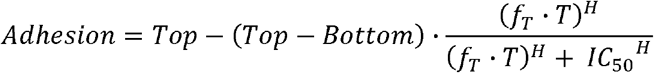

The model was rewritten algebraically to show how one or more orthosteric inhibitors influence the IC50 term.

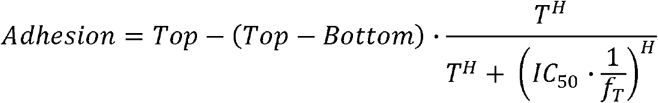

The free fraction of T, f_T_, was expressed as a fold-shift, **α**, of T potency, IC_50_:

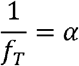

This results in compact expression of the inhibition of TcdB1 effect by one or more inhibitors as a fold-change/fold-shift in the apparent potency of TcdB1:

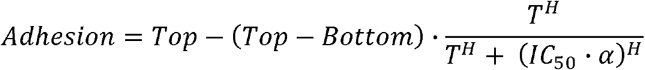

Thus, an experiment can be conducted to estimate effect of TcdB1 alone (yielding an estimate of IC_50_) and in the presence of one or more inhibitors (yielding an estimate of IC_50_·α) to estimate the fold-shift in TcdB1 potency due to the inhibitors (α). While the experimental activities are straightforward, understanding how the inhibitors combine to yield α in a concentration-dependent manner is complex.

We study inhibitor combinations based on their binding patterns to TcdB1. If inhibitors compete for the same site, they follow a competitive binding model, resulting in antagonism under both Loewe additivity and Bliss independence. Inhibitors that do not compete follow an independent binding model, allowing for additivity or synergy in these paradigms.

**Supplemental Table 1** introduces options for the interaction of two or more inhibitors based on their binding epitopes. The potency of each inhibitor is expressed as **K** ^61^, the inhibitor concentration, **C**_i_, that shifts the TcdB1 IC_50_ by exactly 2-fold (α=2):

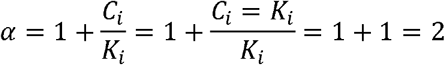

The definition of **K_i_** is like that used in Schild plot analysis where the dose ratio is 2 and “pA_2_” is defined as -log(K), the concentration of inhibitor that results in a dose-ratio (shift) of 2.

During analysis of the data, it became apparent that the linear concentration-potency relationship, 1+C_pp1005_/K_pp1005_, was incompatible with the observed data for pp1005. Specifically, the independent model for three binders (from **Supplemental Table 1**) would be:

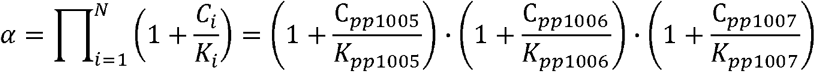

The linear concentration-potency relationship indicates surmountable inhibition of TcdB, as seen with pp1006 and pp1007, where increased TcdB-binding protein counteracts any TcdB concentration. However, pp1005 shows a concentration-dependent limit to inhibition, implying insurmountable inhibition; thus, the model for pp1005 was revised to a saturable Michaelis-Menten type with a Hill coefficient. Extending the independent binding model for all three TcdB-binding proteins yields the final selected synthetic avidity model:

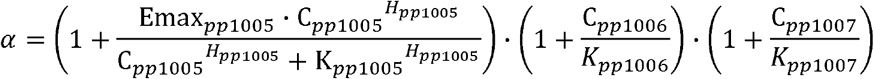

K_i_ values for the TcdB-binding proteins were determined from the totality of the data, which included not only single TcdB-binding proteins but also all double and triple combinations. **Supplemental Table 2** reports the TcdB-binding protein concentration (K_i_) needed to cause a two-fold increase in TcdB IC50 (α=2) for pp1005, pp1006 and pp1007 was 21.44, 54.13, and 69.26 pM, respectively; pp1005 exhibited saturable inhibition with a maximal fold-increase of TcdB IC50 by 45.25-fold (Emax_pp1005_) with a Hill slope coefficient of 1.813 (H_pp1005_).

A comparison to the final independent model was made for the competitive model alternative:

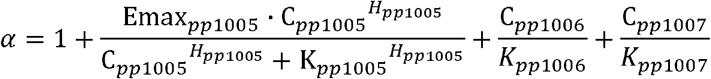

**Supplemental Figure 1** presents model fitting results. The independent model fit the data well and yielded a better goodness-of-fit (by Bayes-Schwartz Information Criteria, or **BIC**) than the competitive model. The competitive model, when fit to TcdB-only (upper left), overpredicted at ∼0.003 pM and underestimated TcdB1 potency. It also overpredicted singleton pp1005 (upper center), underpredicted dual pp1006+pp1007 (lower left), and substantially underpredicted both triple cocktails (lower center and right). Thus, the competitive model failed to account for both highly potent triple combinations and less potent single/dual agents with an additive approach. Extensive searches for interaction permutations and alternative models, including Hill slope for pp1005, were conducted.

**Supplemental Figure 2** demonstrates how the final synthetic avidity model can be used as a simulation tool by examining the response surface of α at four different total TcdB-binding protein concentrations. Simulations were run for a grid of mixtures at these concentrations, and the fold-shift (α) was calculated for each condition. At a total TcdB-binding protein concentration of 0.01 nM (panel A), an almost pure pp1005 mixture is optimal due to pp1005’s superior potency, which is not saturated at this low concentration. As total TcdB-binding protein concentration rises (panels B, C, D), higher proportions of pp1006 and pp1007 become optimal, though a small amount of pp1005 remains beneficial. Analysis of the final synthetic avidity model indicates that maximizing the term for pp1005 to (1+Emax_pp1005_)=(1+45.25) consistently boosts the combined effect of pp1006 and pp1007 by 46.25 fold.

**Supplemental Figure 3** illustrates the application of the final synthetic avidity model as a simulation tool to investigate optimal mixtures across a range of total TcdB-binding protein concentrations, from 0.01 nM to 10 nM. In this context, the optimal mixture is defined as the combination of pp1005, pp1006, and pp1007 that maximizes the fold-shift (α) in TcdB1 potency at the specified total TcdB-binding protein concentration. The resulting optimal mixtures are depicted as a progressive pathway associated with increasing total TcdB-binding protein concentration. At 0.01 nM total TcdB-binding protein, a formulation containing 100% pp1005 (lower left corner) is optimal owing to its superior potency relative to pp1006 and pp1007, consistent with findings presented in Supplemental Figure 2. As the ratio of TcdB-binding protein concentrations to their respective potencies (C_i_/K_i_) exceeds 1.0, the geometric advantage of synthetic avidity yields a mixture of pp1005 and pp1006, visualized as a trajectory toward the upper corner of the diagram. When the effect of pp1005 reaches saturation, its proportion within the mixture declines, yielding a sharp turn in the path towards the centroid of the diagram and onwards to a near-binary mixture of pp1006 and pp1007, supplemented with sufficient pp1005 to achieve maximal, saturated impact.

Analysis of the final synthetic avidity model indicates that the optimal ratio of pp1007:pp1006 is N:1, where N is determined by the ratio K_pp1007_/K_pp1006_ = (0.06926 nM/0.05413 nM), equating to approximately 1.28. Thus, as pp1007 exhibits 1.28-fold lower potency than pp1006 then the maximum product of the two inhibition functions is achieved when pp1007’s concentration is 1.28 times that of pp1006.

**Supplemental Figure 1:**
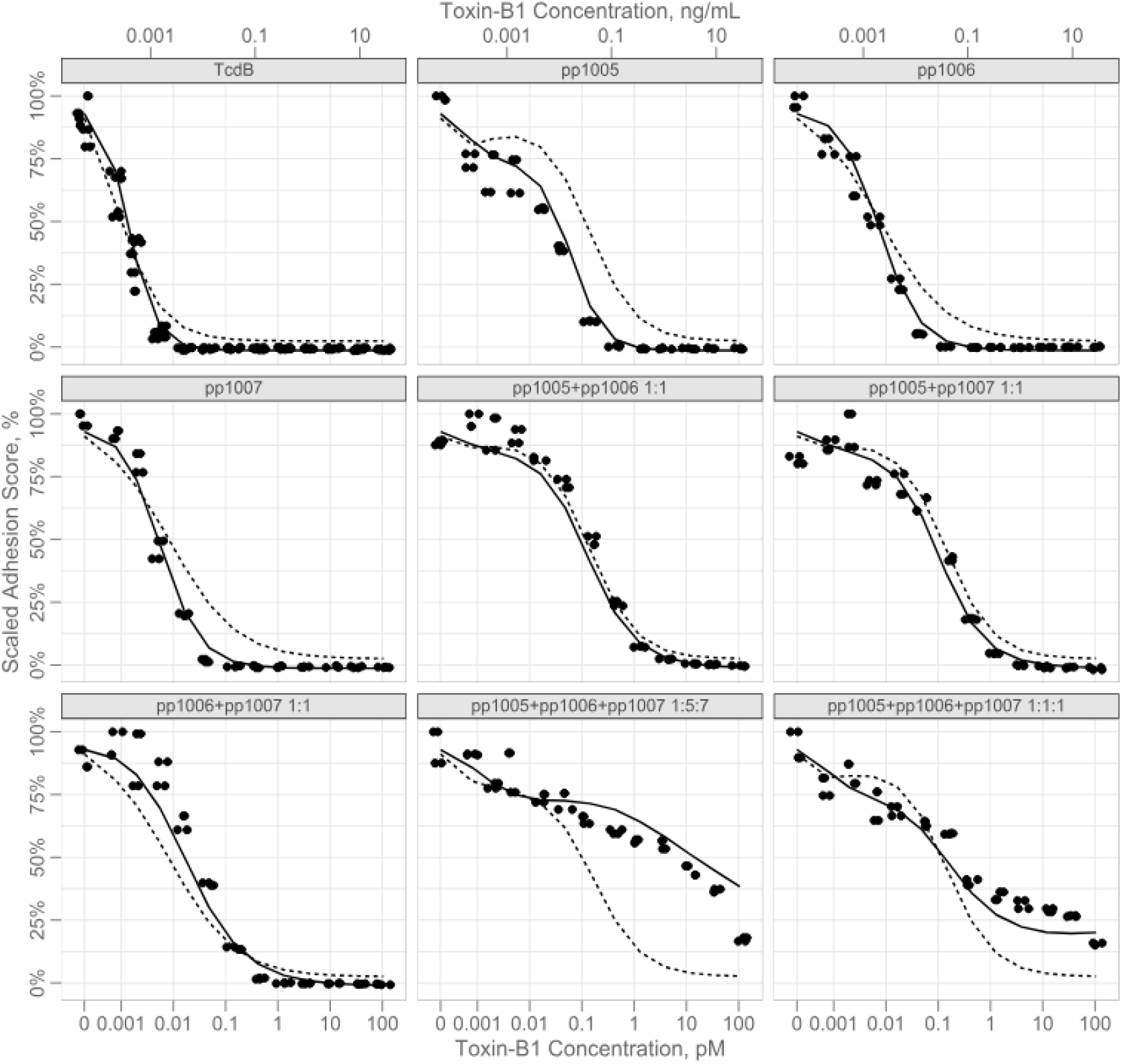
Fitting Results for Independent (Solid Line) and Rejected Alternative Competitive (Dashed Line) Models. The independent model (solid line, BIC=-714) was strongly preferred over the competitive model (dashed line, BIC=-273).

**Supplemental Figure 2.**
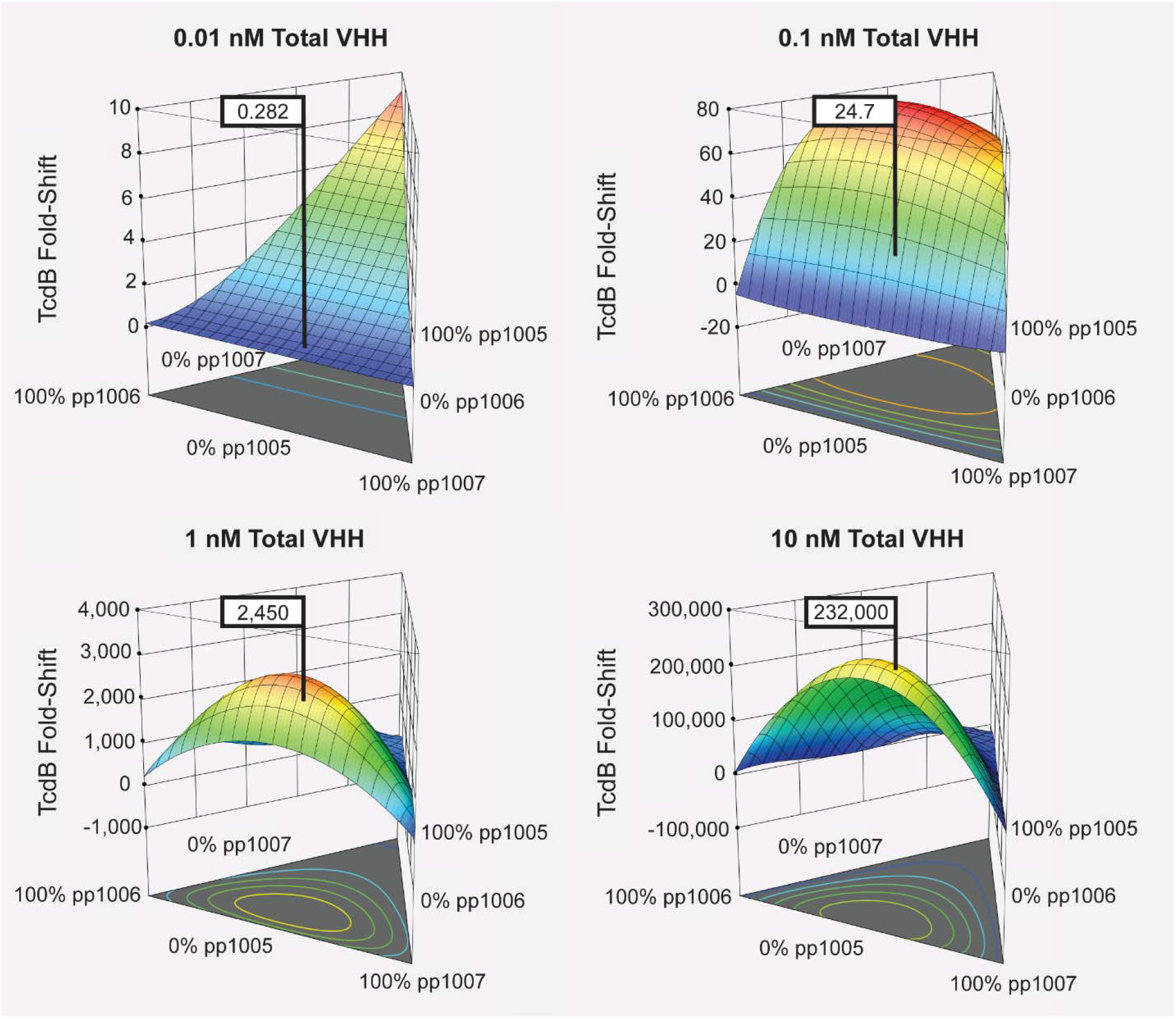
Response Surface Diagrams for Total TcdB-binding Protein Concentrations of 0.01 nM, 0.1 nM, 1 nM and 10 nM. All possible mixtures at these concentrations were simulated, and the fold-shift (α) was calculated for each mixture condition. An example blend of 1:5:7 pp1005:pp1006:pp1007 is highlighted on the response surface.

**Supplemental Figure 3:**
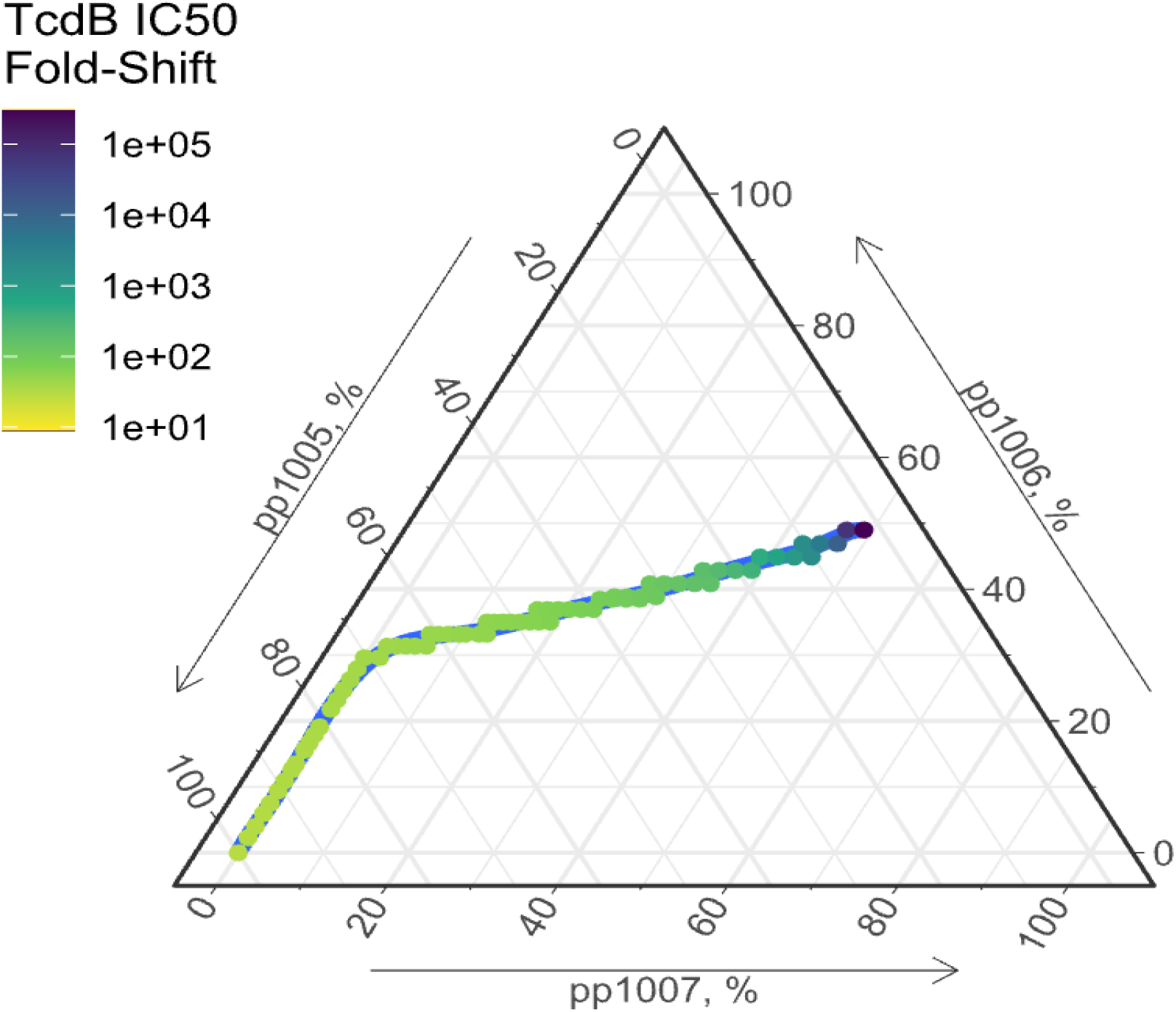
Optimizing the Synthetic Avidity of a Cocktail. Ternary plot showing the optimal mixture of pp1005:pp1006:pp1007 across a total TcdB-binding protein concentration of 0.01 to 10 nM.

**Supplemental Figure 4:**
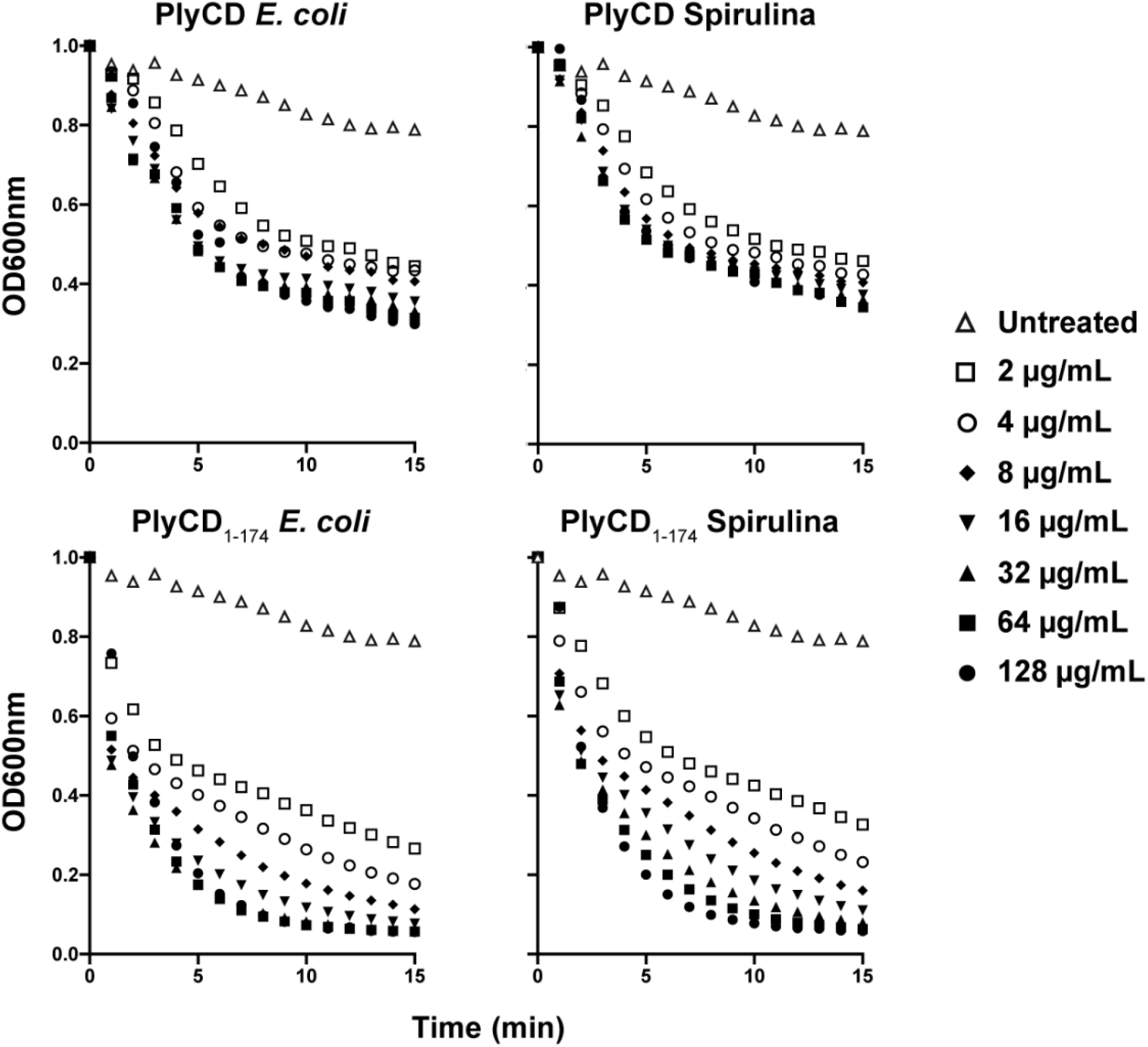
Comparison of PlyCD Lysin Activity Across Expression Systems. Lytic activities of full-length PlyCD (pp1093) and the truncated PlyCD_1–174_ variant (pp1092) expressed in E. coli or spirulina were compared using C. difficile strain ATCC 43255. PlyCD_1–174_ exhibited greater lytic activity than full-length PlyCD, and proteins expressed in spirulina displayed activities comparable to those produced in E. coli.

**Supplemental Figure 5:**
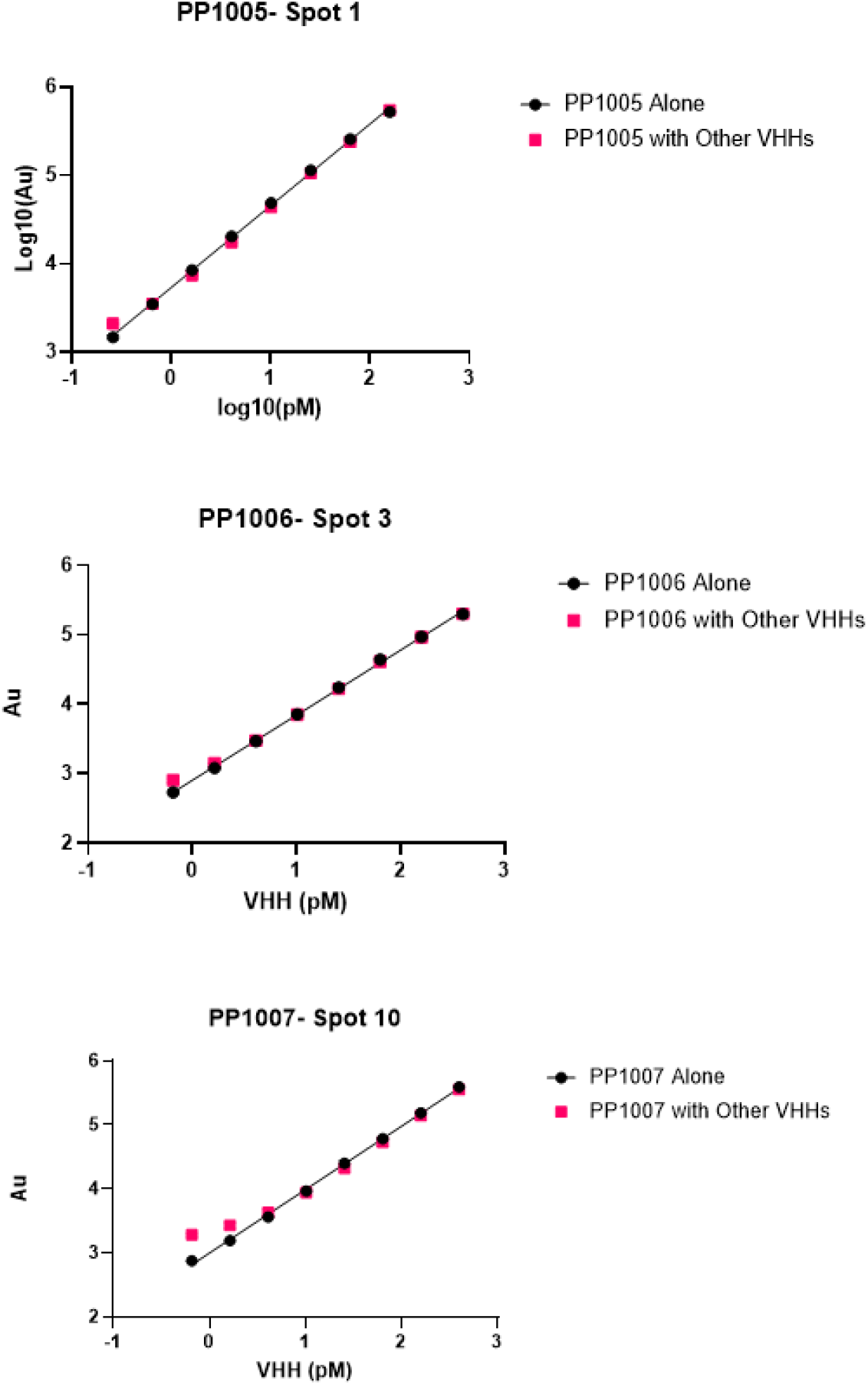
Determination of the Dynamic Range of a Multiplex TcdB-Binding Protein Detection Assay in the Absence of Crosstalk. Each TcdB-binding protein was serially diluted either in buffer alone or in the presence of the maximum tested concentration of each of the other TcdB-binding proteins to define the lower limit of quantification that allowed accurate interpolation from a standard curve generated with the individual TcdB-binding protein. Acceptable assay performance was defined as 80–120% recovery. Black circles denote TcdB-binding proteins diluted in buffer alone, whereas red boxes denote TcdB-binding proteins diluted in the presence of the other TcdB-binding proteins at their maximum concentrations.

**Supplemental Figure 6:**
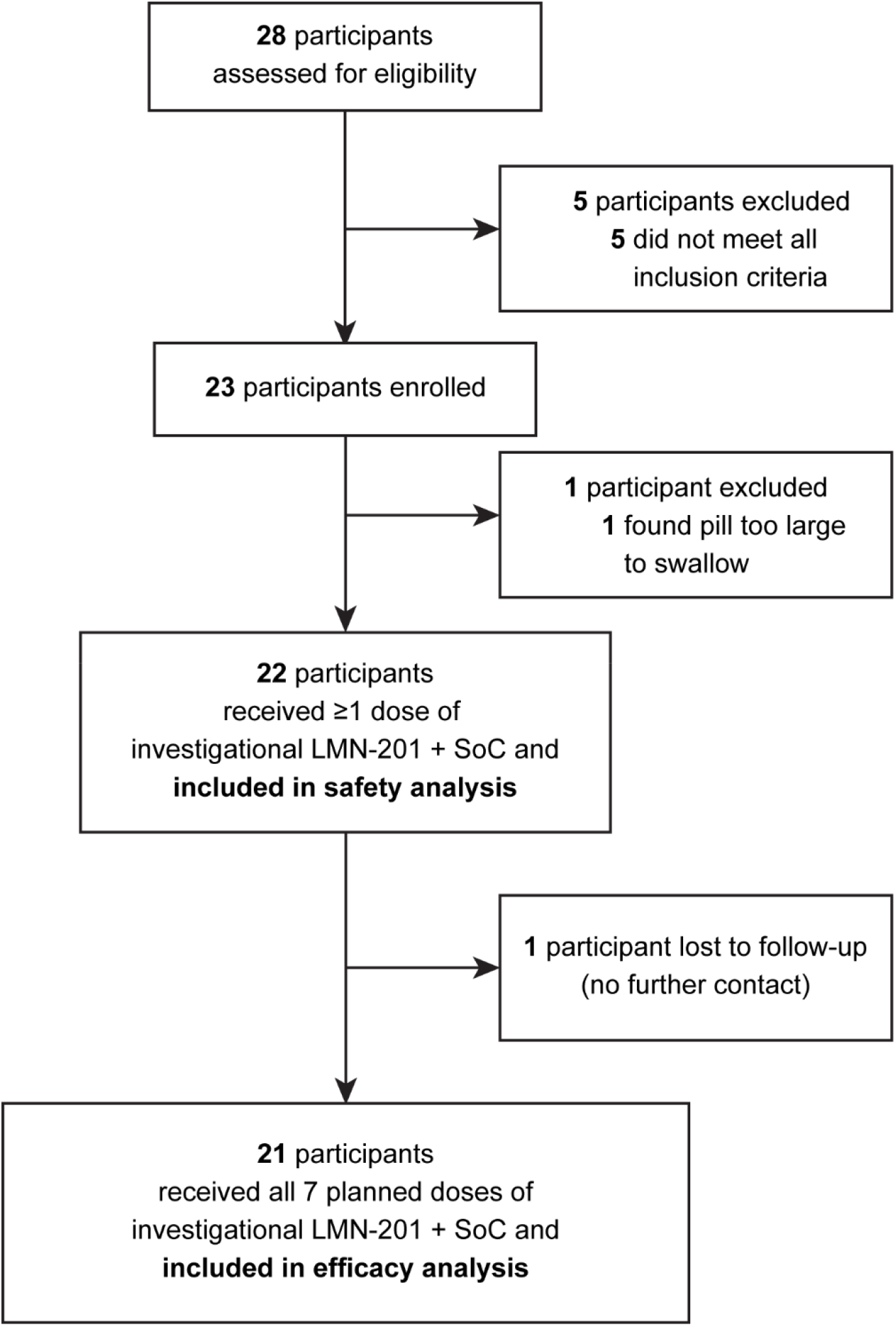
CDI02 clinical trial CONSORT diagram for Part A. SoC = Standard of Care.

**Supplemental Table 1:**
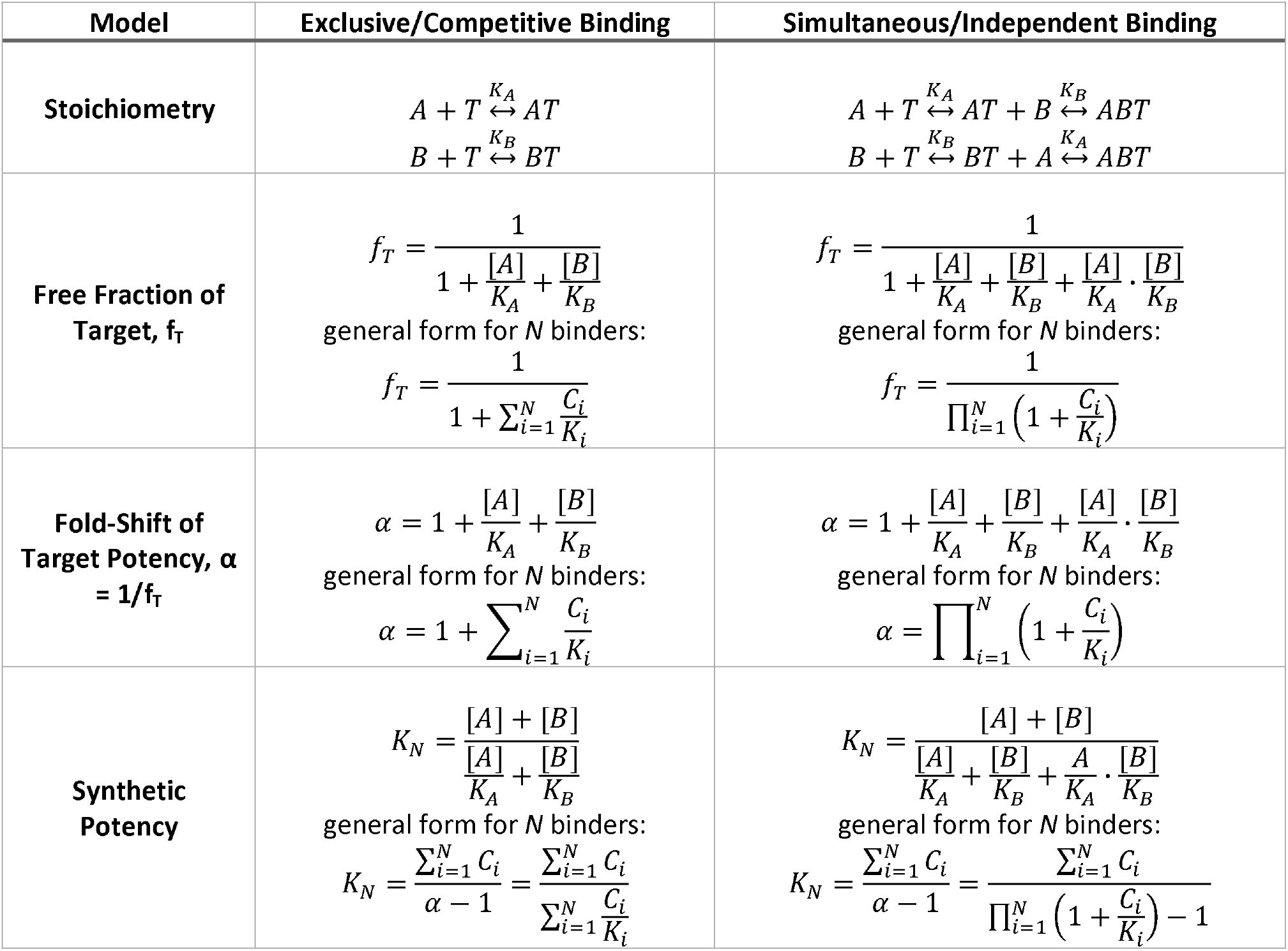
Potential Models Describing Interactions among TcdB-binding Proteins. Differences in stoichiometry, free fraction, fold-shift of target potency and optimal mixture ratios for agents that compete for the same binding site (left column) or do not compete for the same binding site (right column). Stoichiometry reflects association-dissociation of binders (A, B) and target (T) with dissociation equilibrium constants (K_A_, K_B_) to form complexes (AB, BT and ABT).

| **Model** | **Exclusive/Competitive Binding** | **Simultaneous/Independent Binding** |
|---|---|---|
| **Stoichiometry** | $A + T \overset{K_A}{\leftrightarrow} AT$<br>$B + T \overset{K_B}{\leftrightarrow} BT$ | $A + T \overset{K_A}{\leftrightarrow} AT + B \overset{K_B}{\leftrightarrow} ABT$<br>$B + T \overset{K_B}{\leftrightarrow} BT + A \overset{K_A}{\leftrightarrow} ABT$ |
| **Free Fraction of Target, f<sub>T</sub>** | $f_T = \frac{1}{1 + \frac{[A]}{K_A} + \frac{[B]}{K_B}}$<br>general form for *N* binders:<br>$f_T = \frac{1}{1 + \sum_{i=1}^{N} \frac{C_i}{K_i}}$ | $f_T = \frac{1}{1 + \frac{[A]}{K_A} + \frac{[B]}{K_B} + \frac{[A]}{K_A} \cdot \frac{[B]}{K_B}}$<br>general form for *N* binders:<br>$f_T = \frac{1}{\prod_{i=1}^{N} \left(1 + \frac{C_i}{K_i}\right)}$ |
| **Fold-Shift of Target Potency, α = 1/f<sub>T</sub>** | $\alpha = 1 + \frac{[A]}{K_A} + \frac{[B]}{K_B}$<br>general form for *N* binders:<br>$\alpha = 1 + \sum_{i=1}^{N} \frac{C_i}{K_i}$ | $\alpha = 1 + \frac{[A]}{K_A} + \frac{[B]}{K_B} + \frac{[A]}{K_A} \cdot \frac{[B]}{K_B}$<br>general form for *N* binders:<br>$\alpha = \prod_{i=1}^{N} \left(1 + \frac{C_i}{K_i}\right)$ |
| **Synthetic Potency** | $K_N = \frac{[A] + [B]}{\frac{[A]}{K_A} + \frac{[B]}{K_B}}$<br>general form for *N* binders:<br>$K_N = \frac{\sum_{i=1}^{N} C_i}{\alpha - 1} = \frac{\sum_{i=1}^{N} C_i}{\sum_{i=1}^{N} \frac{C_i}{K_i}}$ | $K_N = \frac{[A] + [B]}{\frac{[A]}{K_A} + \frac{[B]}{K_B} + \frac{A}{K_A} \cdot \frac{[B]}{K_B}}$<br>general form for *N* binders:<br>$K_N = \frac{\sum_{i=1}^{N} C_i}{\alpha - 1} = \frac{\sum_{i=1}^{N} C_i}{\prod_{i=1}^{N} \left(1 + \frac{C_i}{K_i}\right) - 1}$ |

**Supplemental Table 2:**
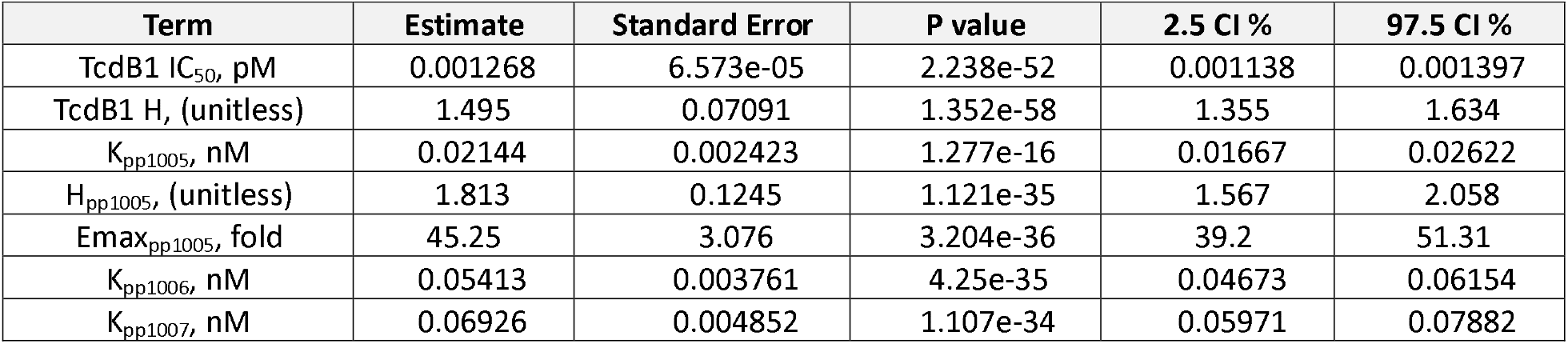
Final Independent Model Parameters.

| **Term** | **Estimate** | **Standard Error** | **P value** | **2.5 CI %** | **97.5 CI %** |
|---|---|---|---|---|---|
| TcdB1 IC<sub>50</sub>, pM | 0.001268 | 6.573e-05 | 2.238e-52 | 0.001138 | 0.001397 |
| TcdB1 H, (unitless) | 1.495 | 0.07091 | 1.352e-58 | 1.355 | 1.634 |
| K<sub>pp1005</sub>, nM | 0.02144 | 0.002423 | 1.277e-16 | 0.01667 | 0.02622 |
| H<sub>pp1005</sub>, (unitless) | 1.813 | 0.1245 | 1.121e-35 | 1.567 | 2.058 |
| Emax<sub>pp1005</sub>, fold | 45.25 | 3.076 | 3.204e-36 | 39.2 | 51.31 |
| K<sub>pp1006</sub>, nM | 0.05413 | 0.003761 | 4.25e-35 | 0.04673 | 0.06154 |
| K<sub>pp1007</sub>, nM | 0.06926 | 0.004852 | 1.107e-34 | 0.05971 | 0.07882 |

**Supplemental Table 3:**
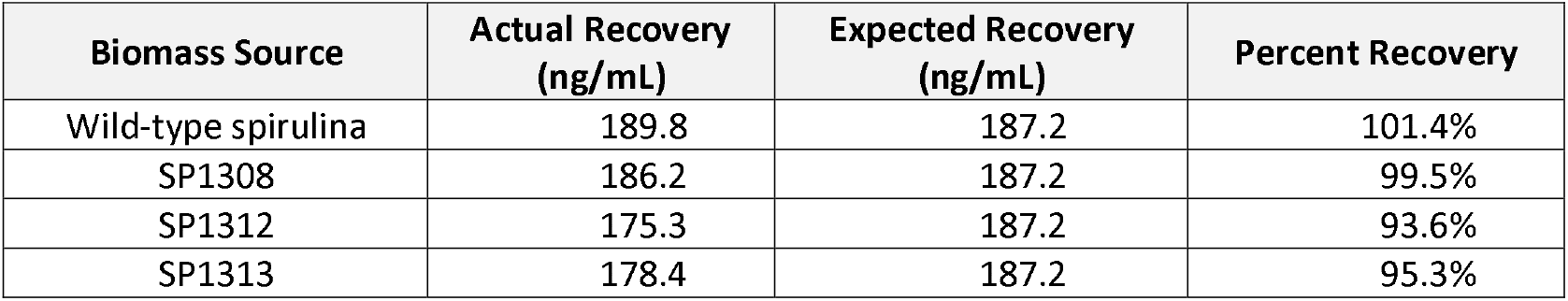
Assessment of Lysin Assay Non-interference by Spirulina-derived TcdB-binding Proteins. Lysates from wild-type spirulina or from strains expressing individual TcdB-binding proteins (SP1308, SP1312 and SP1313) were spiked with equivalent concentrations of pp1092 and analyzed using the turbidity-reduction potency assay. Percent recovery was calculated relative to a standard curve generated in an equivalent concentration of wild-type spirulina matrix. All samples showed percent recovery within acceptable assay variation (80–120%).

### RePreve Trial Supplemental Materials

**Supplemental Table 4:** RePreve Trial Synopsis.

| **Study Description:** | This is a multisite study to evaluate the safety, tolerability, and efficacy of LMN-201 in participants recently diagnosed with CDI who are scheduled to receive or are receiving SoC antibiotic therapy against *C. difficile*. |
|---|---|
| **Population:** | 24 adults with recently diagnosed CDI (clinical symptoms plus positive stool *C. difficile* toxin B immunoassay (see testing algorithm in Section 8.1.10) on stool sample collected in past 10 days) who are scheduled to start receiving SoC antibiotic therapy or have been receiving SoC antibiotic therapy for 10 or fewer days at the time of initial study drug administration. |
| **Inclusion Criteria:** | 1. Male or female, aged 18 years or older.<br>2. Diagnosis of CDI defined as a new or recent history (within past 10 days) of 3 or more bowel movements per day with a loose or watery consistency (Bristol Stool Scale 5, 6, or 7); a positive stool C. difficile toxin B immunoassay (see testing algorithm in Section 8.1.10) on stool collected no more than 10 days before first dose of LMN-201/placebo, and no other likely explanation for diarrhea. NOTE: Diarrhea is not required to be present on the stool sample for CDI diagnosis, on the day of diagnosis, or on the day of enrollment. NOTE: There are no restrictions on prior recurrences.<br>3. Provision of signed and dated informed consent form.<br>4. Scheduled to receive or planning to receive a ≥10-day and ≤28-day course of SoC antibiotic therapy for CDI. If SoC antibiotic therapy has been initiated, participant must have been treated with SoC antibiotic therapy for 10 or fewer days at time of initial study drug administration. For the purposes of this study SoC antibiotic therapy is defined as the receipt of oral fidaxomicin or oral metronidazole or oral vancomycin as described in Section 8.2.6.<br>5. Ability to swallow oral medication (up to size 00 capsules, up to twice per day) and willingness to adhere to the study medication regimen.<br>6. Stated willingness and ability to comply with all study procedures and availability for the duration of the study and investigator believes individual will complete the study.<br>7. Access to and ability to use a mobile smartphone.<br>8. For females of reproductive potential: use of highly effective contraception for at least 4 weeks prior to screening and agreement to use such a method during study participation and for at least 4 weeks after the end of study drug administration.<br>Highly effective contraception is defined as:<br>• Double barrier method: condoms (male or female), diaphragm, or cervical cap with spermicide (foam, gel, film, cream, or suppository)<br>• Surgical sterilization (defined as bilateral oophorectomy, bilateral salpingectomy or bilateral tubal occlusion, performed at least 4 weeks before baseline)<br>• Post-menopausal (defined by amenorrhea for at least 24 months following cessation of all exogenous hormonal treatments)<br>• Combined (estrogen and progesterone containing) hormonal contraception associated with inhibition of ovulation (oral, intravaginal, subdermal, or transdermal).<br>• Progesterone-only hormonal contraception associated with inhibition of ovulation (injectable or implantable)<br>• Intrauterine device (IUD)<br>• Intrauterine hormone-releasing system (IUS)<br>• Sexual intercourse with only vasectomized partners (vasectomy performed at least 6 months before baseline) or<br>• Abstinence is defined as refraining from heterosexual intercourse during the entire period of risk associated with the study treatment. The reliability of sexual abstinence needs to be evaluated in relation to the clinical trial and the preferred and usual lifestyle of the subject.<br>9. For males of reproductive potential: agreement to use condoms or abstinence to ensure effective contraception with partner during study participation and for at least 4 weeks after the end of study drug administration. |
| **Exclusion Criteria:** | 1. Fulminant *C. difficile* colitis (characterized by the development of severe acute inflammation of the colon, associated with systemic toxicity. It is a clinical entity with high mortality that requires intensive medical treatment and early surgery in non-responders).<br>2. Non C. *difficile*-related major GI surgeries, e.g. significant bowel resection, within 3 months of enrollment. NOTE: Does not include appendectomy or cholecystectomy.<br>3. Admitted or expect to be admitted to an intensive care unit.<br>4. Underlying acute (past 3 months) gastrointestinal disorder characterized by diarrhea. Examples include but are not limited to short bowel syndrome, dumping syndrome following gastrectomy, enteric parasitic infection, viral enteritis, bacterial enteritis (salmonella, shigella, enterotoxigenic *E. coli*, etc.).<br>5. Underlying active chronic (past year) gastrointestinal disorder characterized by diarrhea. Examples include but are not limited to chronic ulcerative colitis, Crohn's disease, celiac sprue, pancreatic insufficiency, inflammatory bowel disease (IBD).<br>6. Current or previous treatment in past 3 months with any therapy likely to influence the outcome of this study, including but not limited to the following:<br>a. Bezlotoxumab (Zinplava, Merck & Co.), or another antibody against *C. difficile* toxin(s)<br>b. *C. difficile* vaccine<br>c. SER-109 (Seres Therapeutics)<br>d. CP101 (Finch Therapeutics)<br>e. VE303 (Vedanta Therapeutics)<br>f. Rebyota (Ferring Pharmaceuticals)<br>g. Other fecal microbiota transplant<br>h. Current therapy with oral exchange resins<br>7. Known current history of significantly compromised immune system such as:<br>a. Known history of human immunodeficiency virus (HIV) infection and CD4 < 200 cells/mm<sup>3</sup> within 6 months of start of enrollment.<br>b. Severe neutropenia with neutrophil count < 500 cells/mL.<br>c. Concurrent intensive induction chemotherapy, radiotherapy, or biologic treatment for active malignancy.<br>8. Pregnancy, anticipated pregnancy within the study timeline, or breastfeeding.<br>9. Known allergy or sensitivity to corn, spirulina, or other study drug constituents.<br>10. Psychiatric illness that, in the opinion of the Investigator, would affect compliance with medications, study drug, or follow-up.<br>11. Terminal illness with limited life expectancy of less than 24 weeks.<br>12. Concurrent medical risks with clinically significant co-morbid disease such that, in the opinion of the investigator, the participant should not be enrolled.<br>13. Any other condition that, in the opinion of the investigator, would jeopardize the safety or rights of the individual, would make it unlikely for the individual to complete the study, or would confound the results of the study.<br>• Note: Use of probiotics and other food supplements (e.g., yogurt, kefir, kimchi, etc.) are not exclusionary.<br>• Note: Use of anti-diarrheals, opiates, or other anti-motility agents is not exclusionary. |
| **Enrolling Sites:** | Outpatient clinic, hospital, long-term care facility |
| **Participant Duration:** | Participant duration from screening through last safety follow-up is approximately 27 weeks.<br>Study phases are broken out over a 27-week study period as follows:<br>• Treatment Phase with LMN-201 twice daily dosing for 7 days overlapping with SoC antibiotic therapy<br>• Observation Phase with no study drug, observation for safety for adverse events, serious adverse events, and medically attended adverse events for 4 weeks<br>• Follow-up Phase for safety for serious adverse events and medically attended adverse events for 22 weeks, starting after the completion of the Observation Phase |

**Supplemental Table 5:** Comparison of Enrollment Criteria between MODIFY I/II and RePreve Trial Part A.

| **Inclusion Criteria** |  |  |  |
|---|---|---|---|
| Age | Adults only: “18 years of age or older.” *(nejmoa1602615_protocol.pdf p.25)* | Adults only: “aged 18 years or older.” *(CDI02 v3.0 p.26)* | Comparable (adult-only). |
| CDI diagnosis requirement | CDI defined by both: (a) “3 or more loose stools in 24 or fewer hours” and (b) “a positive stool test for toxigenic C. difficile.” Also, the stool sample with positive result must be collected “within 7 days prior to the infusion.” *(nejmoa1602615_protocol.pdf p.25–26)* | CDI defined by: “3 or more bowel movements per day with a loose/watery consistency (Bristol 5–7)”; “positive stool C. difficile toxin B immunoassay… on stool collected no more than 10 days before first dose”; and “no other likely explanation.” Also notes: diarrhea not required to be present on diagnosis/enrollment day; no restrictions on prior occurrences/recurrences. *(CDI02 v3.0 p.26)* | RePreve is explicitly more permissive on symptom timing (diarrhea not required at enrollment) and includes an explicit “no other explanation” clause. MODIFY is more “classic”: diarrhea + positive toxigenic test near infusion. |
| Diagnostic Lab Assays | MODIFY-I/II included stool culture of the organism, a series of PCR tests, and a list of toxin tests. *“Positive stool test for toxigenic C. difficile or toxin(s).”* Acceptable tests included: Cell Culture Cytotoxin Assays, OR Stool Culture with Toxigenic Strain Typing, OR Stool Culture with Toxin Detection from *C. difficile* isolates, OR one of the following commercial assays: | RePreve requires a positive toxin B immunoassay in addition to any other tests that might be run. This is to ensure as much as possible that the patient has active infection. *Positive stool test for C. difficile toxin B immunoassay on a stool sample collected not more than 10 days prior to the date of first dose of study drug.* | MODIFY I/II diagnostic criteria were broader than RePreve, and could include multiple methods of identifying CDI. This was reflective of diagnostic guidelines in the mid-2010s when the protocol was conducted.<br>As a result, the RePreve trial is less likely to include individuals who have been colonized with *C. difficile*. |
|  | **The acceptable tests for the studies are:** |  |  |
|  | **Manufacturer** | **Assay Name** | **CDI02 or MODIFY** |
|  | Becton Dickinson | BD GeneOhm | MODIFY only |
|  | Biomerieux | VIDAS C. difficile Toxins A and B | CDI02 and MODIFY |
|  | Cepheid | GeneXpert | MODIFY only |
|  | Medical Chemical Corp. | GastroTect | MODIFY only |
|  | Meridian | Premier | CDI02 and MODIFY |
|  |  | Illumigene | MODIFY only |
|  |  | Immunocard Toxins A and B | CDI02 and MODIFY |
|  | Oxoid/Remel | Xpect C. difficile Toxin A/B | CDI02 and MODIFY |
|  |  | ProSpecT A and B | CDI02 and MODIFY |
|  | Prodesse | ProGastro CD | MODIFY only |
|  | TechLab (or Inverness Medical) | Tox A/B QuikChek | MODIFY only |
|  |  | C. diff QuikChek Complete | MODIFY only |
|  |  | C. difficile Tox A/B II | MODIFY only |
|  | Alere / TechLab | C. DIFF QUIK CHEK COMPLETE Immunoassay | CDI02 only |
|  |  | C. difficile Tox A/B II | CDI02 only |
|  |  | TOX A/B QUIK CHEK | CDI02 only |
| **Exclusion Criteria** |  |  |  |
| Fulminant/severe CDI / ICU | Does *not* appear as a single “fulminant CDI” exclusion in the exclusion list; instead severe disease is handled in definitions/subgroups (e.g., Zar severity score includes ICU treatment as 2 points). *(nejmoa1602615_protocol.pdf p.67–68, p.28)* | Explicit exclusions: “Fulminant *C. difficile* colitis… high mortality”; “Admitted or expect to be admitted to an ICU.” *(CDI02 v3.0 p.27)* | RePreve is more restrictive—may exclude the sickest CDI patients who were eligible in MODIFY (or at least not explicitly excluded). |
| Chronic diarrheal illnesses / IBD | Excludes “active chronic diarrheal illness… ulcerative colitis or Crohn’s disease.” *(nejmoa1602615_protocol.pdf p.26)* | Excludes both: “acute (past 3 months) GI disorder characterized by diarrhea” and “active chronic (past year) GI disorder… ulcerative colitis, Crohn’s, celiac… IBD.” *(CDI02 v3.0 p.27)* | RePreve is more detailed and time-bound (acute vs chronic windows). Both exclude IBD and other potential confounders. |
| Major GI surgery | Excludes “planned surgery for CDI within 24 hours.” *(nejmoa1602615_protocol.pdf p.26)* | Excludes “major GI surgeries… significant bowel resection within 3 months.” *(CDI02 v3.0 p.27)* | RePreve focuses on recent non-CDI bowel surgery; MODIFY focuses on imminent CDI surgery. |
| Prior biologics / microbiome products targeting CDI | Excludes certain prior/concomitant therapies (e.g., “fidaxomicin within 14 days”; “cholestyramine, rifaximin, nitazoxanide, or fidaxomicin within 14 days”; “immune globulin within 6 months”; planned antiperistaltics and certain probiotics such as *S. boulardii*). *(nejmoa1602615_protocol.pdf p.26–27)* | Explicitly excludes prior 3-month exposure to many CDI-directed interventions: “Bezlotoxumab… C. difficile vaccine… SER-109… CP101… VE303… Rebyota… other FMT… oral exchange resins.” *(CDI02 v3.0 p.28)* Also: probiotics/food supplements and anti-motility agents are not exclusionary. *(CDI02 v3.0 p.28)* | This is a major divergence: RePreve is explicitly designed to avoid confounding from recent anti-CDI biologics/microbiome products, while MODIFY restricts certain meds and planned agents but is less “product-specific” reflecting therapies available at the time. |
| Immunocompromise | Not a blanket exclusion; rather, “compromised immunity” is defined and used for subgroup analyses. *(nejmoa1602615_protocol.pdf p.68)* | Explicit exclusion: “significantly compromised immune system” examples include HIV CD4 <200, severe neutropenia, intensive chemo/radiation/biologics for active malignancy. *(CDI02 v3.0 p.28)* | RePreve is more conservative on immunosuppression. This may reduce generalizability but potentially lowers safety risk and outcome noise. |
| Pregnancy and breastfeeding | Excludes positive pregnancy test and breastfeeding, plus donation/contraception restrictions. *(nejmoa1602615_protocol.pdf p.26–27)* | Excludes “pregnancy… or breastfeeding.” *(CDI02 v3.0 p.28)* | Both standard; MODIFY includes more detailed contraception/donation language. |

## Author Contributions

**Conceptualization:** Michael Dodds, Michael Tasch, Nhi Khuong, Brian Finrow, and James M. Roberts Data generation and curation: Hui Zhao, Alex Pollock, Mesfin Gewe, Anissa Martinez, Kristie Keeney, Benjamin W. Jester, Melanie Hutton, Hannah Tabakh, Mia Zhang, Mark Heinnickel, Troy Paddock, Stacey Ertel, Kanchan Aggarwal, Caroline R. Amendola, Colin Brady, Nathaniel R. Sanjaya, Madison Kahwaty, Kendra Cruickshank, Sarah Struyvenberg, Jason Dang, Tracy Saveria, Chelsea Shanitta, Yena Park, David Doughty, Jackie Poirot, Caitlin Pensak, Thomas Adame, Jeremy Ferrara, Aaron Read, Dalton Heil, Spencer Shibuya, James “Chip” Henderson, David Fletcher, Taylor Charron, Kristjan Sigmar, Eva Aw, René Ruiz, Karen Yoshino, James Lo, Dorothy Strong, Lauren Goetsch, Caitlin Gamble, Yasi Zhong, Ryan Antolock, Cosette Loper, Gabby Guzman, Sophie Gist, Steven J. Mileto, Ryan Heselpoth, Michael Tasch, Carl J. Mason, Nhi Khuong

**Formal analysis:** Hui Zhao, Michael Dodds

**Funding acquisition:** Brian Finrow, Nhi Khuong, James M. Roberts

**Investigation:** Hui Zhao, Michael Dodds, Alex Pollock, Mesfin Gewe, Melanie Hutton, David Saunders, George B. McDonald, Michael Tasch, Summer Radler, Carl J. Mason, Craig A. Behnke, Nhi Khuong, and James M. Roberts

**Methodology:** Hui Zhao, Michael Dodds, Alex Pollock, Mesfin Gewe, Hannah Tabakh, Mark Heinnickel, Troy Paddock, Michael Tasch, David Doughty, Carl J. Mason

**Project administration:** Michael Dodds, Sarah Shephard, Asa Davis, Sanika Khare, Ashley Zambrano, Michael Tasch, Carl J. Mason, Craig A. Behnke, Nhi Khuong, and James M. Roberts

**Supervision:** Hui Zhao, Michael Dodds, Alex Pollock, Mesfin Gewe, Caitlin Gamble, Asa Davis, David Saunders, Dena Lyras, Vincent Fischetti, Michael Tasch, Summer Radler, Carl J. Mason, Craig A. Behnke, Nhi Khuong, Brian Finrow, and James M. Roberts

**Visualization:** Hui Zhao, Michael Dodds, Alex Pollock, Mesfin Gewe, Kristie Keeney, Benjamin W. Jester, Hannah Tabakh, Mark Heinnickel, Samuel T. Robinson

**Writing – original draft:** Hui Zhao, Michael Dodds, Alex Pollock, Mesfin Gewe, Kristie Keeney, Benjamin W. Jester, Hannah Tabakh, Mark Heinnickel, Carl J. Mason, Nhi Khuong, and James M. Roberts

**Writing – review & editing:** Hui Zhao, Michael Dodds, Brian Finrow, James M. Roberts, Samuel T. Robinson

## Declaration of generative AI and AI-assisted technologies in the writing process

During the preparation of this work, the authors used ChatGPT in order to ensure clarity and consistency within the text. After using this tool/service, the authors reviewed and edited the content as needed and take full responsibility for the content of the published article.

## Acknowledgements

This work was supported by the Assistant Secretary of Defense for Health Affairs endorsed by the Department of Defense, in the amount of $16,288,194 through the Peer Reviewed Medical Research Program under Award No. HT9425-23-1-0959. Opinions, interpretations, conclusions and recommendations are those of the authors and are not necessarily endorsed by the Assistant Secretary of Defense for Health Affairs or the Department of Defense. Lumen Bioscience, Inc. has utilized the non-clinical and pre-clinical services program offered by the National Institute of Allergy and Infectious Diseases to facilitate the hamster study. Lumen Bioscience, Inc. also used internal funding for the completion of some of these studies and this manuscript.

We thank Charles Shoemaker, George McDonald, and Fred Cross for help, discussion, and advice. We thank Meagan James and other members of the Lyras laboratory for assistance with the mouse studies.

For the hamster challenge studies, Lumen Bioscience has utilized the non-clinical and pre-clinical services program offered by the National Institute of Allergy and Infectious Diseases, especially support from Sangun Lee, Christian Gonzalez, and Ryan Ranallo.

We thank the members of the Lumen Bioscience research and development teams for their scientific input, technical assistance, and project support throughout this work. We are especially grateful to colleagues involved in strain engineering, protein production, strain growth optimization, analytical characterization, and assay development for their contributions to the advancement of this program.

We acknowledge the Lumen Bioscience Production team for their efforts in manufacturing clinical trial materials, and the Quality Assurance and Quality Control teams for their essential support in release testing, quality oversight, and preparation of materials for clinical studies.

We thank the Clinical team for their contributions to the planning, coordination, execution, and oversight of the clinical trial activities associated with this program.

We acknowledge the Project Management team for their leadership, coordination, and cross-functional support throughout the program. We also thank the IT and Data Infrastructure team for their support with data management, digital infrastructure, and operational systems that facilitated the execution and coordination of this work.

We thank colleagues who contributed to manuscript preparation, editing, and coordination throughout the writing process.

We thank the Lumen Bioscience Laboratory Operations team for their general laboratory support, coordination, and assistance in maintaining research infrastructure and day- to-day operations that enabled this work.

We also acknowledge the Business Development and Legal teams for their support with external partnerships, contractual, compliance, and intellectual property matters related to this research.

